# Centromere scission drives chromosome shuffling and reproductive isolation

**DOI:** 10.1101/809848

**Authors:** Vikas Yadav, Sheng Sun, Marco A. Coelho, Joseph Heitman

## Abstract

A fundamental characteristic of eukaryotic organisms is the generation of genetic variation via sexual reproduction. Conversely, significant large-scale genome structure variations could hamper sexual reproduction, causing reproductive isolation and promote speciation. The underlying processes behind large-scale genome rearrangements are not well understood and include chromosome translocations involving centromeres. Recent genomic studies in the *Cryptococcus* species complex revealed that chromosome translocations generated via centromere recombination have reshaped the genomes of different species. In this study, multiple DNA double-strand breaks (DSBs) were generated via the CRISPR/Cas9 system at centromere-specific retrotransposons in the human fungal pathogen *Cryptococcus neoformans*. The resulting DSBs were repaired in a complex manner, leading to the formation of multiple inter-chromosomal rearrangements and new telomeres, similar to chromothripsis-like events. The newly generated strains harboring chromosome translocations exhibited normal vegetative growth but failed to undergo successful sexual reproduction with the parental wild-type strain. One of these strains failed to produce any spores, while another produced ∼3% viable progeny. The germinated progeny exhibited aneuploidy for multiple chromosomes and showed improved fertility with both parents. All chromosome translocation events were accompanied without any detectable change in gene sequences and thus, suggest that chromosomal translocations alone may play an underappreciated role in the onset of reproductive isolation and speciation.

## Introduction

Chromosomes are prone to undergo several rearrangement events, including fusion, fission, deletion, and segmental duplication. In some cases, one chromosome segment is exchanged with another to generate chromosomal translocations. Such exchanges between homologs are regularly observed during meiosis when homologous chromosomes exchange arms via meiotic recombination (1). Chromosome rearrangements can also occur during mitosis, but in a less well-regulated manner, and sometimes as a result of disease conditions like cancer (2, 3). Additionally, rearrangements can also occur within a single chromosome. As a result, chromosome rearrangements during mitosis can cause mutations, gene disruption, copy number variations, as well as alter the expression of genes near the breakpoints (4). Cancer cells show a high level of chromosome rearrangements as compared to healthy cells, and this contributes to critical pathological conditions observed in these cells, such as activation of oncogenes (5, 6).

Chromosomal translocations are initiated by double-strand breaks (DSBs) in DNA (7). Rearrangements involving a single DSB are mainly repaired by the invasion of the broken DNA molecule into a homologous DNA molecule, in a process termed homologous recombination (HR) (6, 8). This invasion can lead to the exchange of DNA between the two molecules of DNA, leading to reciprocal crossover or gene conversion (2). These types of rearrangements occur during meiosis and are regulated to give rise to an error-free repaired sequence. Other types of chromosomal translocation involve two or more DNA DSB sites, which are then fused in an error-prone mechanism known as non-homologous end joining (NHEJ) (9). The two sites can be present on either the same chromosome or different chromosomes. Repair of two DSBs present on the same chromosome can result in the deletion of the intervening sequence or inversion if the sequence is rejoined in reverse orientation (7). On the other hand, the fusion of DSBs from two different chromosomes can result in chromosomal translocation. In some instances, other repair pathways like microhomology-mediated end joining (MMEJ) or alternative End Joining (alt-EJ) also participate in the repair of DSB ends (9, 10).

The occurrence of multiple DSBs in mitotically growing cells at the same time is rare but occurs at a relatively higher frequency in cancer cells. Such events could occur in a genome due to replication defects or exogenous factors such as ionizing radiation or chemotherapeutic agents (11). Multiple DSBs also occur naturally during processes like V(D)J recombination (12). The occurrence of multiple DSBs can also be induced in the micronuclei of cancer cells (13). Micronuclei are small nuclei harboring one or a few chromosomes and are generated as a result of a mitotic failure. These small nuclei act as hotspots of chromosome fragmentation, where multiple DSB sites are almost simultaneously generated and subsequently rejoined in a random order, a process known as chromothripsis. Previous reports have suggested that the occurrence of multiple DSBs alters HR pathways leading to NHEJ or alt-EJ mediated repair (14, 15). Because both NHEJ and alt-EJ are error-prone, they lead to a significant increase in mutation at the repair junctions and also randomly join broken fragments.

Apart from generating mutations and gene disruptions, chromosome translocations can also result in reproductive isolation during meiosis and facilitate speciation (16). The presence of multiple rearrangements between the two homologous chromosomes from the parents leads to failures in chromosome pairing during meiosis or crossovers that result in loss of essential genes (17). Meiosis that is defective in this way will result in the production of progeny with abnormal genome content. In fungi, the parental nuclei fuse and undergo meiosis before sporulation that gives rise to progeny. Thus, defects in meiosis lead to the production of spores with abnormal or incomplete genetic compositions rendering them inviable. *Cryptococcus neoformans* is a basidiomycete fungus that largely infects immunocompromised humans causing Cryptococcal meningoencephalitis (18–20). *C. neoformans* harbors a 19 Mb genome with 14 chromosomes (21). While most of the genome is devoid of repeat regions, centromeres in *C. neoformans* are rich in a set of LTR-retrotransposons (251 full-length and truncated copies of 6 elements) that are shared across multiple centromeres (22, 23). *C. neoformans* also has a meiotic cycle making it a suitable model organism to study meiosis and sexual development (20, 24). It exhibits two mating types, α and **a**, defined by the *MAT*α and *MAT***a** alleles at the single mating-type locus (25, 26).

In this study, we exploited the genomic features of *C. neoformans* to study the impact of chromosome translocations on reproductive isolation. First, retrotransposons present in centromeres were targeted with CRISPR, generating multiple DSBs simultaneously. Next, the presence of chromosome rearrangements was screened by Pulsed-Field Gel Electrophoresis (PFGE), and isolates with multiple chromosomal translocations were identified. The genomes of these strains were assembled based on long-read nanopore sequencing to characterize the chromosome rearrangements. Although the strains with new karyotypes did not exhibit growth defects as compared to the wild-type, the impact of chromosomal rearrangements had a profound effect on sexual reproduction. These findings demonstrate that *C. neoformans* can tolerate multiple chromosomal translocations, but that such large-scale changes can cause reproductive isolation, and promote incipient speciation.

## Results

### Simultaneous breaks at multiple centromeres lead to chromosome shuffling

Centromeres in *C. neoformans* were previously identified and shown to possess multiple retrotransposons named Tcn1-Tcn6 (21, 22). These elements are present only in centromeres, albeit distributed randomly and non-uniformly across all 14 centromeres. We specifically targeted the Tcn2 element using a guide RNA (gRNA) that would cleave nine centromeres, generating a total of 18 DSBs (Fig. 1A). We hypothesized that these DSBs would be repaired to generate chromosomal translocations (Fig. 1B).

**Fig. 1.**
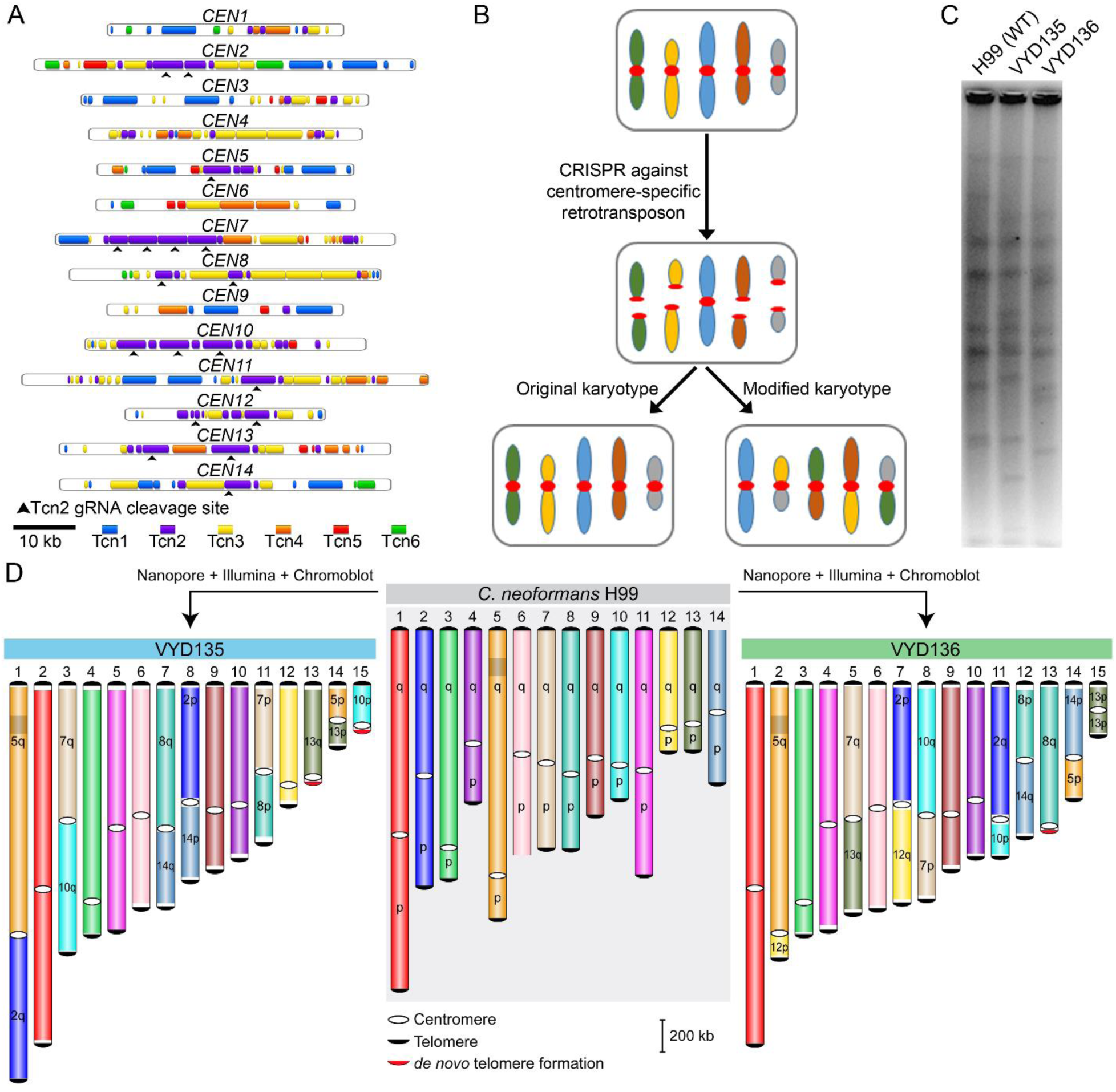
Centromere-specific DSBs mediated by CRISPR lead to chromosome rearrangements. **(A)** Centromere maps showing the distribution of retrotransposons (Tcn1-Tcn6) in the centromeres of wild-type strain H99 of *C. neoformans*. **(B)** An outline depicting the model for achieving multiple chromosome translocations in *C. neoformans*. **(C)** PFGE revealed many differences in the karyotype of VYD135 and VYD136 as compared with wild-type, H99. **(D)** Chromosome maps for VYD135 and VYD136 compared to the H99 genome revealed multiple chromosome translocations in these strains. Chromosomes are colored with H99 chromosomes as reference. ‘q’ represents the longer arm while ‘p’ represents the shorter arm according to the wild-type chromosome configuration.

After the transformation of wild-type cells with three DNA fragments, each constitutively expressing a dominant selection marker (G418), Cas9, and the gRNA, colonies were obtained and screened via pulsed-field gel electrophoresis (PFGE) (Fig. S1). We found karyotypic variation for at least one chromosome in three out of twelve colonies screened (Fig. S2A). Of the three, two (VYD135 and VYD136) harbored more than three changes in chromosome banding pattern as compared to the wild-type strain *C. neoformans* H99 (Fig. 1C and Fig. S2A). To characterize these chromosomal alterations, we sequenced the genomes of these two strains using Oxford Nanopore sequencing and were able to assemble them into 17 contigs each (Fig. S2B).

A synteny block comparison of these genomes with that of the wild-type revealed the presence of multiple translocations. However, some of these contigs were broken with the centromere at one of the ends. Additionally, Illumina sequencing of these strains revealed a near euploid genome for both of these strains (Fig. S2C) except for the smaller arm of chromosome 13 in VYD136 that was present in two copies. To resolve the status of incompletely assembled contigs, Southern blots of PFGE-separated chromosomes (chromoblots) followed by hybridization with chromosome-specific probes were conducted for multiple chromosomes (Fig. S3). Contigs validated to be arms of the same chromosome by this approach were fused manually with a 50-bp sequence gap to generate a chromosome-size scaffold. The presence of telomere repeat sequences at both the ends of these scaffolds suggests that these scaffolds represent full-length chromosomes. The integrity of each genome assembly was further verified by mapping the respective nanopore reads to the genome assembly. Thus, using nanopore and Illumina sequencing, and chromoblot analysis, we were able to assemble the genomes of isolates VYD135 and VYD136 to the chromosome level (Fig. 1D). This analysis also revealed that the duplicated arm of chromosome 13 of isolate VYD136 exists as an isochromosome with two broken centromeres fused with each other. Overall, these results show that multiple breaks at centromeres can lead to karyotype shuffling in *C. neoformans* that is tolerated by the organism.

### Centromere breaks generate new telomeres and increase the number of chromosomes

No species in the *Cryptococcus* species complex has been observed to harbor chromosomes smaller than 500 kb. However, genome-level assemblies of VYD135 and VYD136 revealed the presence of 2 to 3 chromosomes that are shorter than the shortest naturally occurring chromosome of wild-type H99 (chromosome 13 of 757 kb length). Bands of the expected size for these chromosomes were observed in PFGE, indicating that these chromosomes are *bona fide* and not a result of assembly error (Fig. 2A). In addition to these small chromosomes, a few novel features were observed for the genomes of VYD135 and VYD136. Three of the new chromosomes had generated *de novo* telomere sequences on one of the ends, next to the broken centromeres (chromosomes 13 and 15 of VYD135; chromosome 13 of VYD136) (Fig. 2B, and Fig. S4A). While in two cases, the telomere repeat sequences were present next to the Tcn2 elements, in the third multiple copies of the Cas9 sequence were found to be inserted between the Tcn2 element and telomere repeat sequences (Chr15 of VYD135). Our analysis of these regions did not reveal any common motif as the most likely target for the *de novo* telomere addition. While one (Chr13 of VYD135) of the *de novo* additions was preceded by a sequence similar to that of a telomere repeat (Table S1), the other two telomeres did not show such a feature. This suggests that the addition of telomere repeat sequences may randomly occur on broken chromosome ends.

**Fig. 2.**
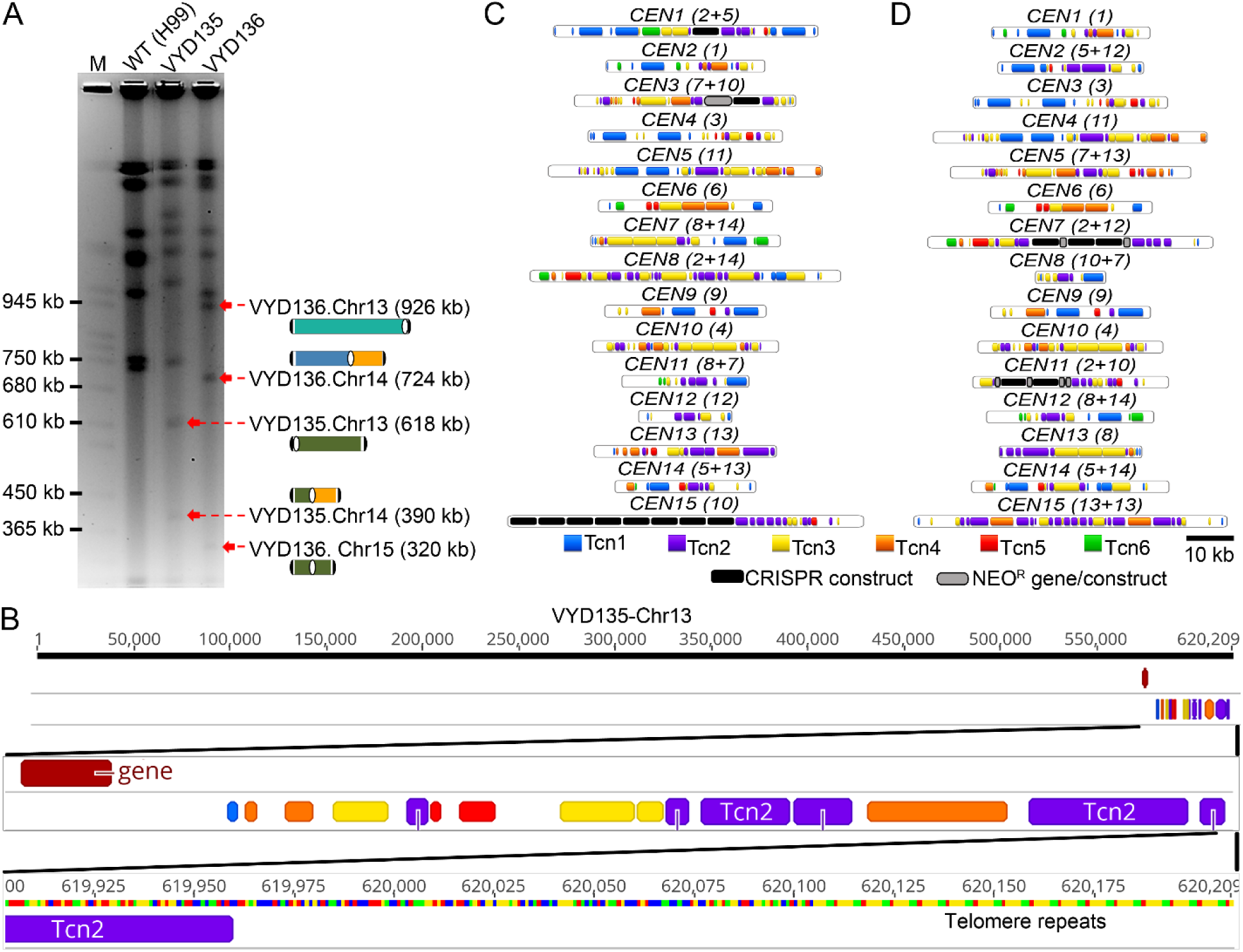
Sequence analysis of new centromeres and telomeres. **(A)** PFGE EtBr staining showing bands for newly generated short chromosomes and new telocentric chromosomes. M represents *S. cerevisiae* chromosomes. **(B)** Chromosome map of one of the newly formed telomeres showing the presence of telomere sequence repeats next to Tcn2 elements present in the centromere. **(C-D)** Maps showing the distribution of retrotransposons, along with the integration of foreign sequences from CRISPR/Cas9 and the neomycin resistance gene (NEO^R^), in the centromeres of VYD135 (C) and VYD136 (D). Numbers in brackets next to *CEN* numbers represent the wild-type *CEN* numbers that rearranged to form new centromeres.

Centromere locations in these two genome assemblies were defined based on synteny with centromere flanking regions (Table S2). Specifically, BLASTn analysis with centromere-flanking ORF sequences as query sequences were used to identify the syntenic regions defining the boundary for centromeres. Next, Tcn elements (Tcn1-6) were mapped across centromeres of the new strains, VYD135 and VYD136. Surprisingly, foreign DNA elements, such as Cas9 and neomycin resistance gene sequence, were found to have integrated into multiple centromeres (Fig. 2C and 2D). Both of these sequences were introduced as linear DNA molecules during the CRISPR transformation experiment. In some cases, Cas9 and the neomycin gene sequences were present in multiple copies and in a random order/orientation. Further analysis revealed that these elements were present at the junction where two parental centromeres fused with each other. This result suggests that these foreign sequences may have assisted in joining the broken centromeres to form hybrid centromeres. Previous studies in *S. cerevisiae* observed similar events where the Ty1 retrotransposon cDNA sequence was found to be present at DNA break sites (27, 28).

A comparison of centromere length between H99, VYD135, and VYD136 revealed the presence of some significantly shorter and longer centromeres (16-83 kb versus 31-64 kb in H99) in both of these new strains (Fig. S4B and Table S2). However, the variation in centromere length did not seem to confer a visible growth defect indicating no change in centromere function due to a decrease or increase of length (Fig. S5A). These strains also did not show any difference when tested for various stress conditions such as temperature, fluconazole, or DNA damaging agent like Phleomycin (Fig. S5A). When grown with another strain expressing NAT resistant gene, to test competitive fitness, both VYD135 and VYD136 did not show any significant difference as compared to wild-type H99 (Fig. S5B). Additionally, these two strains do not exhibit any difference in virulence in the *Galleria* model of infection (29) (Fig. S5C). Both VYD135 and VYD136 lead to lethal infection of *Galleria* with the same efficiency as the wild-type H99α and KN99**a** isolates. Taken together, these results suggest that the changes in chromosome structure do not grossly affect mitotic fitness or infectivity of *C. neoformans*.

### Chromosome shuffling is driven by the Cas9 induced breaks

Isolates VYD135 and VYD136 show inter-chromosomal rearrangements between seven and eight chromosomes, respectively. The rearrangements in the two strains are not identical, although they do involve the same set of chromosomes, suggesting that both of these strains underwent recombination via different routes (Fig. 3A and 3B). Next, the breakpoints in each of the centromeres were defined by mapping nanopore reads to the wild-type genome. One caveat is that long reads from nanopore sequencing might not align precisely with the parental centromere sequence due to the loss of regions of the original sequence. Many reads were found to be mapped to a single location next to the gRNA cleavage site in almost all of the centromeres that underwent recombination (Fig. 3C and Fig. S6). In a few cases, the reads did not map to the cleavage site flanking sequences suggesting deletions occurred during recombination. This loss of sequence is prominent in centromeres with three or more cleavage sites (*CEN10* in Fig. 3C). This mapping pattern suggests these non-essential, small fragments were lost, and their loss does not compromise centromere function. Interestingly, among the chromosomes that contain Tcn2 elements and could be targeted by Cas9 with the Tcn2-specific gRNA, chromosome 11 was not found to be involved in recombination in either of the two strains, even though *CEN11* is predicted to be cleaved once. It is possible that the gRNA cleavage site prediction for *CEN11* could be a result of an incorrect genome assembly or sequence error. Overall, this analysis supports that recombination was initiated by the Cas9 DSBs. Also, that centromeres lacking Tcn2 elements were not involved in recombination reflects the specificity of Cas9 and the repair machinery.

**Fig. 3.**
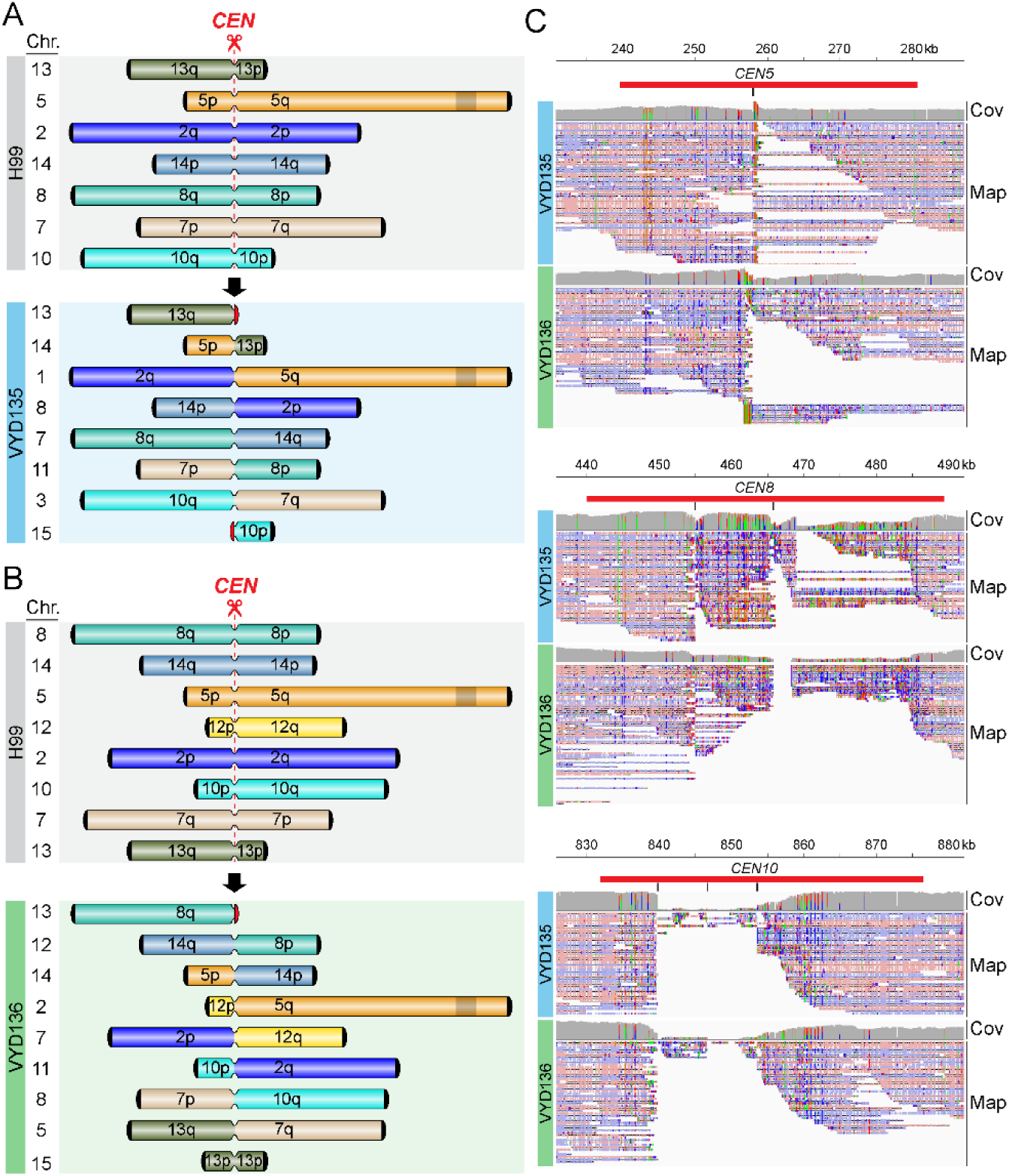
Chromosome rearrangements are mediated via the gRNA cleavage site. **(A-B)** Simplified outline maps depicting the chromosomes that underwent translocation in VYD135 (A) and VYD136 (B). Black semicircles, telomeres; red semicircles, de novo telomeres, narrow bands, centromeres; shaded box on chromosome 5, *MAT* locus. **(C)** Nanopore reads mapping to the wild-type H99 genome revealed the sites of DSB formation or repair junctions at centromeres. The reads either converge on a single site (*CEN5*, *CEN8*) or exhibit sequence gaps between sites (*CEN10*) marking the location of junctions. Red bars indicate the centromeres whereas the black vertical lines mark the site of gRNA cleavage. Cov, Coverage of nanopore reads; Map, Mapping of nanopore reads.

### Multiple types of repair machinery were involved in the DSB repair process

Next, the new centromere sequences were compared with the original centromeres in a pairwise fashion to understand the repair process (Fig. 4 and Fig. S7). For this purpose, we utilized our newly generated assembly of the wild-type strain, which showed better coverage for centromere sequences (See Materials and Methods for details). This analysis provides evidence that both NHEJ and HR pathways participated in the repair of these broken ends. Insertion of the *CAS9* sequence and the *NEO* gene sequence conferring G418 resistance seems to be the result of DSB repair via NHEJ in all cases as there was no additional sequence added between the ends of the breaks (Fig. 4B, 4F, and Fig. S7D, S7E). The presence of these DNA fragments in multiple copies, due to transformation, might have facilitated their insertion. Additionally, *CEN12* of VYD136 results from a single fusion event between wild-type *CEN8* and *CEN14* after the DNA breaks (Fig. S7A). For *CEN14* of VYD135, a sequence of 2.7 kb aligned with both parental centromeres suggesting that the hybrid centromere formed as a result of homologous recombination (Fig. 4A). *CEN8* of VYD135 exhibited evidence for repair via multiple mechanisms, including NHEJ, HR, and invasion into multiple different centromeres (Fig. 4C). Because all of the sequences involved have a Tcn2 sequence at the ends, it is not possible to infer the order of these events.

**Fig. 4.**
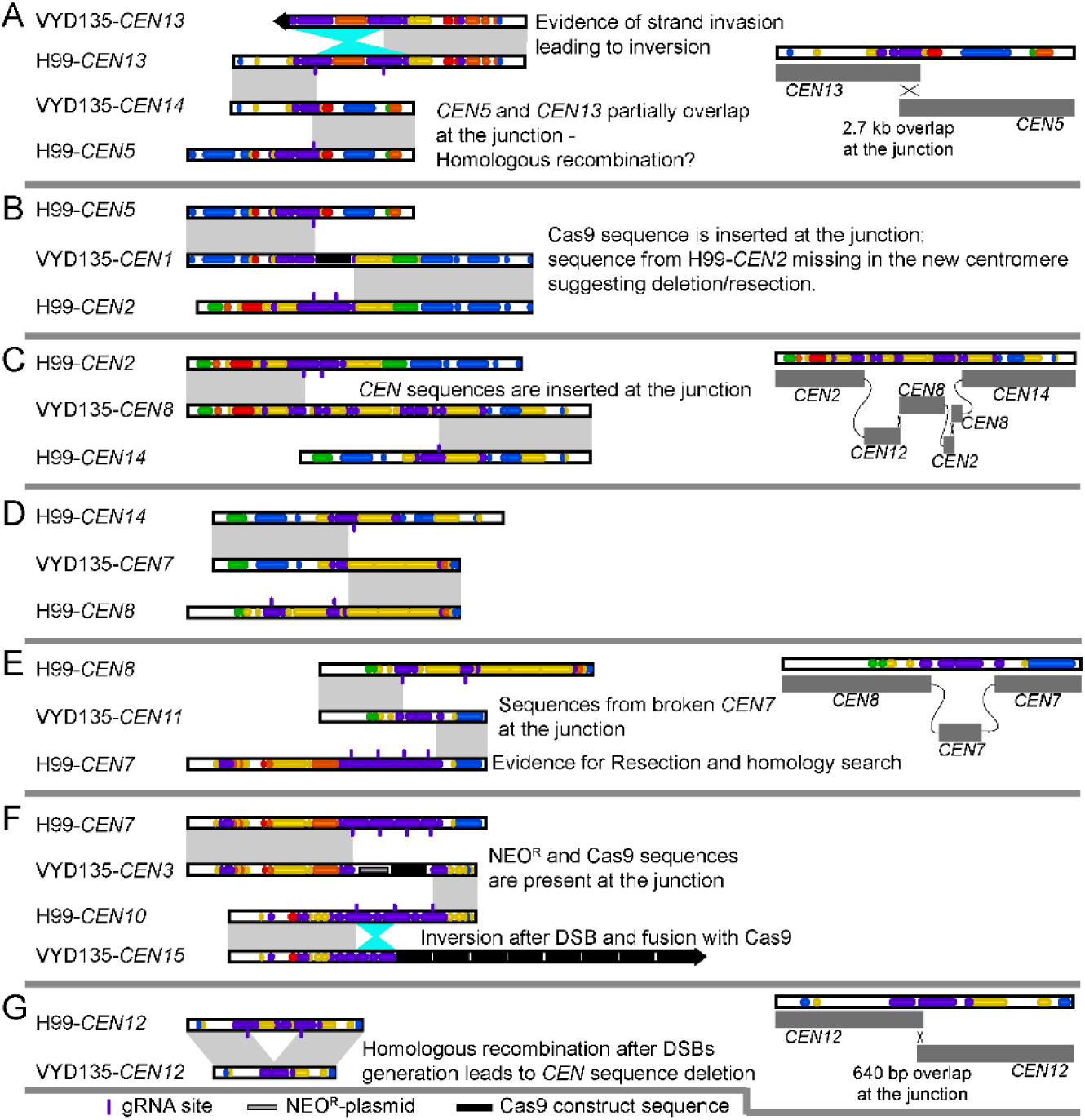
Synteny analysis of rearranged centromeres with wild-type H99 centromeres reveals complex rearrangements. **(A-G)** A pair-wise comparison of newly generated and wild-type centromeres revealed that translocations are mediated by double-strand breaks (DSBs) generated via CRISPR. Centromere specific events are described in the individual panels. Grey shades represent direct synteny, while the cyan shade represents inversion events. In the cases that are shown in detail, the cross represents the evidence for homologous recombination, whereas the connecting grey lines represent joining events marking non-homologous end-joining. VYD135-*CEN7* was generated after artificial fusion of two contigs and hence was not analyzed in detail.

A comparison of wild-type *CEN7* and VYD135 *CEN11* shows evidence of resection beyond the DSB sites (Fig. 4E). Resection was probably followed by strand invasion into broken pieces of *CEN7* that were released due to multiple DSBs, and a final fusion event with *CEN8* leading to the formation of hybrid centromeres. Similarly, *CEN13* of VYD136 seems to have arisen as a result of resection followed by invasion into multiple centromeres before adding telomere sequences at the end (Fig. S7A). In addition to these multiple recombination events at these junctions, we also observed inversion events for sequences that were released due to multiple DSBs in a single centromere (*CEN15* of VYD135) (Fig. 4F). On the other hand, the inversion in VYD135 *CEN13* has signatures of invasion into another intact copy of *CEN13*, resulting in the inversion (Fig. 4A).

Apart from inter-chromosome recombination, we also observed intra-chromosomal recombination in *CEN12* of VYD135. Wild type *CEN12* has two cleavage sites separated by a 10 kb sequence. In VYD135, *CEN12* is smaller due to the absence of the 10 kb sequence, and two flanking sequences were joined with an overlap of 640 bp (Fig. 4G). This event also led to a reduction in centromere length for *CEN12*, shortening it significantly (21 kb versus 31 kb in the wild type). Combined, these results suggest that both HR and NHEJ processes repair DSBs at the centromeres. Given the high level of identity shared between Tcn2 elements present among multiple centromeres, other plausible routes to these rearrangements are possible.

### Strains with chromosomal rearrangements fail to undergo normal meiosis

Chromosome shuffled strains, VYD135 and VYD136, exhibit multiple chromosomal translocations compared to the wild-type karyotype, as described above. We hypothesized that this would lead to incompatibility during meiosis and defects in producing viable spores. To study this, the two shuffled strains were crossed with the wild-type strain, KN99**a**, which is congenic with the parental strain H99α. During *Cryptococcus* sexual reproduction, cells of two opposite mating types fuse and then produce hyphae, basidia, and long chains of spores over a two-week course of incubation (26). In the basidium, the two parental haploid nuclei fuse and undergo meiosis producing four haploid nuclei. These meiotic products subsequently undergo repeated rounds of mitosis that produce four spore chains from the surface of the basidium by budding. A pairing defect during meiosis would compromise the segregation of chromosomes, giving rise to progeny with incomplete or imbalanced genomes.

After two weeks of mating, intact spore chains were observed by light microscopy in the H99α x KN99**a** wild-type cross. On the other hand, the VYD135α x KN99**a** cross formed basidia but showed almost no sporulation, whereas the VYD136α x KN99**a** cross was largely indistinguishable from wild-type and did not show a prominent sporulation defect (Fig. 5A). Meiotic structures were further examined by scanning electron microscopy (SEM). Abundant spores were produced in the H99α x KN99**a** cross, whereas no spore chains were observed in the VYD135 x KN99**a** cross (Fig. 5B). The VYD136 x KN99**a** cross showed defective sporulation, where some basidia formed fewer spores while others had none (Fig. 5B). Next, the germination rate (equivalent to spore viability) of spores from the H99α x KN99**a** and VYD136 x KN99**a** crosses was assessed. The spore germination rate for the VYD136 x KN99**a** cross was only 3% compared to wild-type with 80 to 90% spore germination (30, 31). The lack of sporulation in the VYD135 x KN99**a** cross and the reduced spore germination rate for the VYD136 x KN99**a** cross provide evidence that these strains fail to undergo normal meiosis. The translocations in VYD135 involve seven chromosomes, and in VYD136 eight chromosomes are involved, yet these isolates show very different phenotypes during meiosis. We hypothesize that this observed difference in the sporulation phenotype may be attributable to different chromosome configurations observed in these two strains. Specifically, different configurations could lead to differences in meiotic pairing and could cause more severe meiotic defects in one compared to the other. The failure to undergo meiosis will lead to no or defective sporulation.

**Fig. 5.**
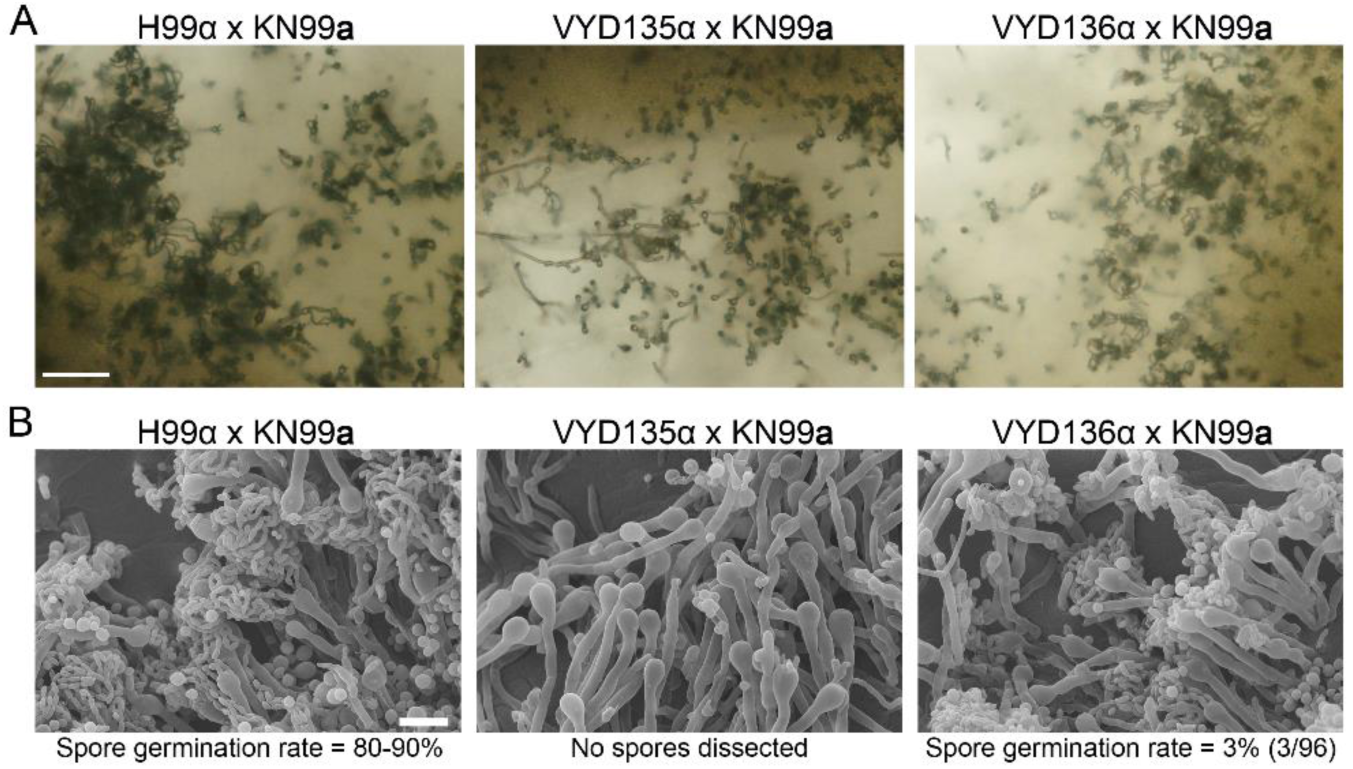
Chromosome shuffled isolates exhibit defects in sexual reproduction. **(A)** Light microscopy images showing hyphae, basidia, and spore chains in crosses between *MAT***a** wild-type strain KN99**a** and *MAT*α wild-type (H99) and rearranged strains (VYD135 and VYD136). Scale bar, 100 µm. **(B)** Scanning Electron Microscope (SEM) images depicting a complete or partial sporulation defect when wild-type KN99**a** was crossed with VYD135 and VYD136, respectively. Scale bar, 10 µm.

### VYD136 backcross progeny are aneuploid and exhibit improved sexual reproduction

Three viable F1 progeny were isolated from the cross of VYD136 with KN99**a**. These F1 progeny (P1, P2, P3) were characterized to further understand the impact of chromosomal translocations on meiosis by assessing mating type and ploidy (Fig. 6A and 6B). Flow cytometry analysis revealed that two of the progeny (P1 and P2) are aneuploid. To investigate this further, the relative chromosome copy number for the three progeny was determined from read counts after Illumina and nanopore sequencing, revealing that they are aneuploid for multiple chromosomes (Fig. 6C). P1 and P2 are both aneuploid for parts of chromosomes 5, 12, 13, and 14, whereas P2 is also aneuploid for the entire chromosome 6. The third progeny (P3) also showed aneuploidy but only for the shorter arms of chromosomes 12 and 13.

**Fig. 6.**
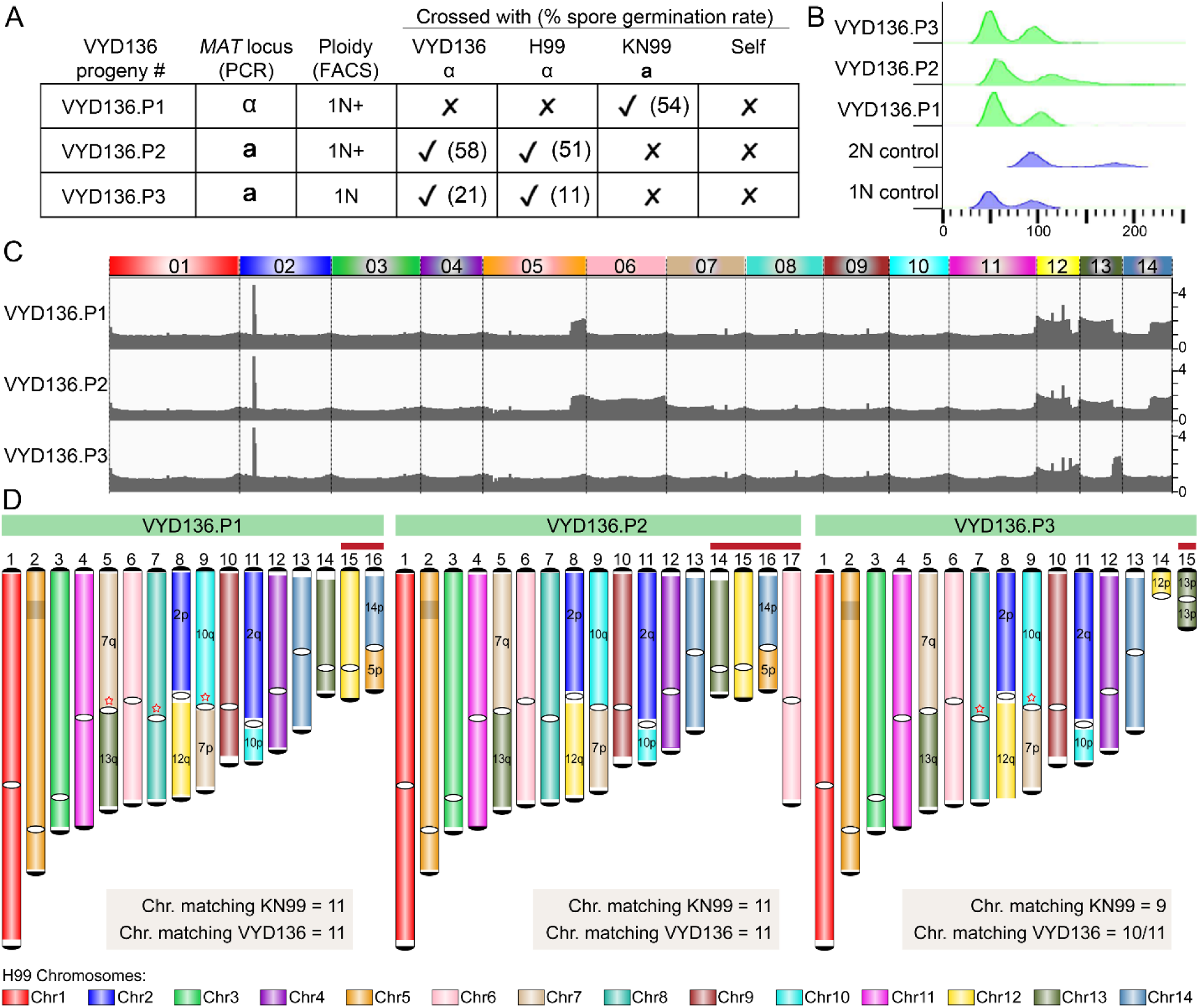
VYD136 progeny are aneuploid and exhibit mixed karyotypes. **(A)** Analysis of the mating-type locus and the mating efficiency of three progeny of VYD136 with either parent. The numbers in brackets represent spore germination rates of respective crosses. **(B)** Flow cytometry profiles of wild-type haploid, diploid, and three progeny of VYD136. **(C)** Illumina sequencing data mapped to the wild-type H99 genome revealed aneuploidy for multiple chromosomes in the three VYD136 progeny. **(D)** Karyotypes for three progeny showing synteny as compared to the wild-type H99 genome. The red stars represent breaks that were fused later based on synteny and ploidy. The chromosomes shown with red bar on top were not assembled *de novo* but represent possible chromosomes configuration based on Illumina and nanopore sequencing analysis. Contigs 14 in VYD136.P3 could not be resolved into their chromosome configuration. ‘q’ represents longer arm while ‘p’ represents shorter arm according to the wild-type chromosome configuration.

These three F1 progeny were backcrossed to both parents (VYD136 and KN99**a**) and the wild-type H99α. All three progeny mated with strains of opposite mating type, as expected. Surprisingly, spore dissections from these crosses revealed a much higher spore germination rate for the three progeny (11 to 58%) as compared to their parental cross (3% germination rate) (Fig. 6A). P1 and P2 exhibited a much higher (>50%) germination rate for all crosses as compared to P3, which showed 11 and 21% germination rates. The segmental aneuploidy of these isolates may explain their higher germination rate compared to the parent VYD136.

The presence of copy number changes for only one arm of most chromosomes suggested these progeny may have a mixed karyotype. To address this, genomes for the three progeny were assembled using nanopore sequencing. Due to aneuploidy, these genomes were not assembled completely and harbored multiple breaks (Fig. S8A and S8B). However, based on their ploidy profiles, mapping of nanopore reads, as well as synteny comparison with both parents, their genome configurations were largely resolved (Fig. 6D). The final karyotype shows that most chromosomes match either with the wild-type or the VYD136 chromosome profile (Fig. 6D and Fig. S8A). Based on this, we propose that segmental aneuploidy may contribute to overcome the reproductive barrier that might otherwise arise due to multiple changes in chromosome configuration. Genetic transmission via an aneuploid intermediate may yield a transitional population that is more compatible with the ancestral population and slowly diverges to give rise to a new species. On the other hand, a new species might also arise because of a single event of multiple translocations occurring throughout the genome. This process is only one of several factors contributing to reproductive isolation and could play a greater or lesser role dependent upon the species distribution and mode of reproduction.

## Discussion

Chromosome rearrangements shape the genome in multiple ways by affecting gene positions, causing mutations, and compromising recombination during meiosis. In this study, chromosomal rearrangements were generated by severing centromeres in a human fungal pathogen, *C. neoformans*. The centromeres in this species are rich in retrotransposons that are common among multiple centromeres (22). CRISPR/Cas9 targeting of retrotransposons cleaved eight centromeres simultaneously in each case. This approach does not affect gene-rich regions, thus avoiding the risks involved with the loss or mutation of genes. At the same time, this allowed us to study a) how multiple breaks in the genome are tolerated, b) how DSBs in heterochromatic regions such as centromeres are repaired, and c) how the resulting chromosomal rearrangements impact sexual reproduction. Centromere-mediated chromosomal translocation events have been observed in the *Cryptococcus* species complex (21, 32) (Fig. 7A-F). Genome comparisons between non-pathogenic *C. amylolentus* and pathogenic species, such as *C. neoformans*, show at least six centromere-mediated translocations (32). Previous studies have suggested that retrotransposons, including centromeric elements, are actively transcribed and can undergo transposition in *Cryptococcus* species (22, 33, 34). Thus, it is possible that these elements generate DNA breaks during their insertion in to the centromeres, which can then catalyze the occurrence of chromosome rearrangements. Our findings suggest these translocations could have driven speciation in this species complex, in addition to other factors.

**Fig. 7.**
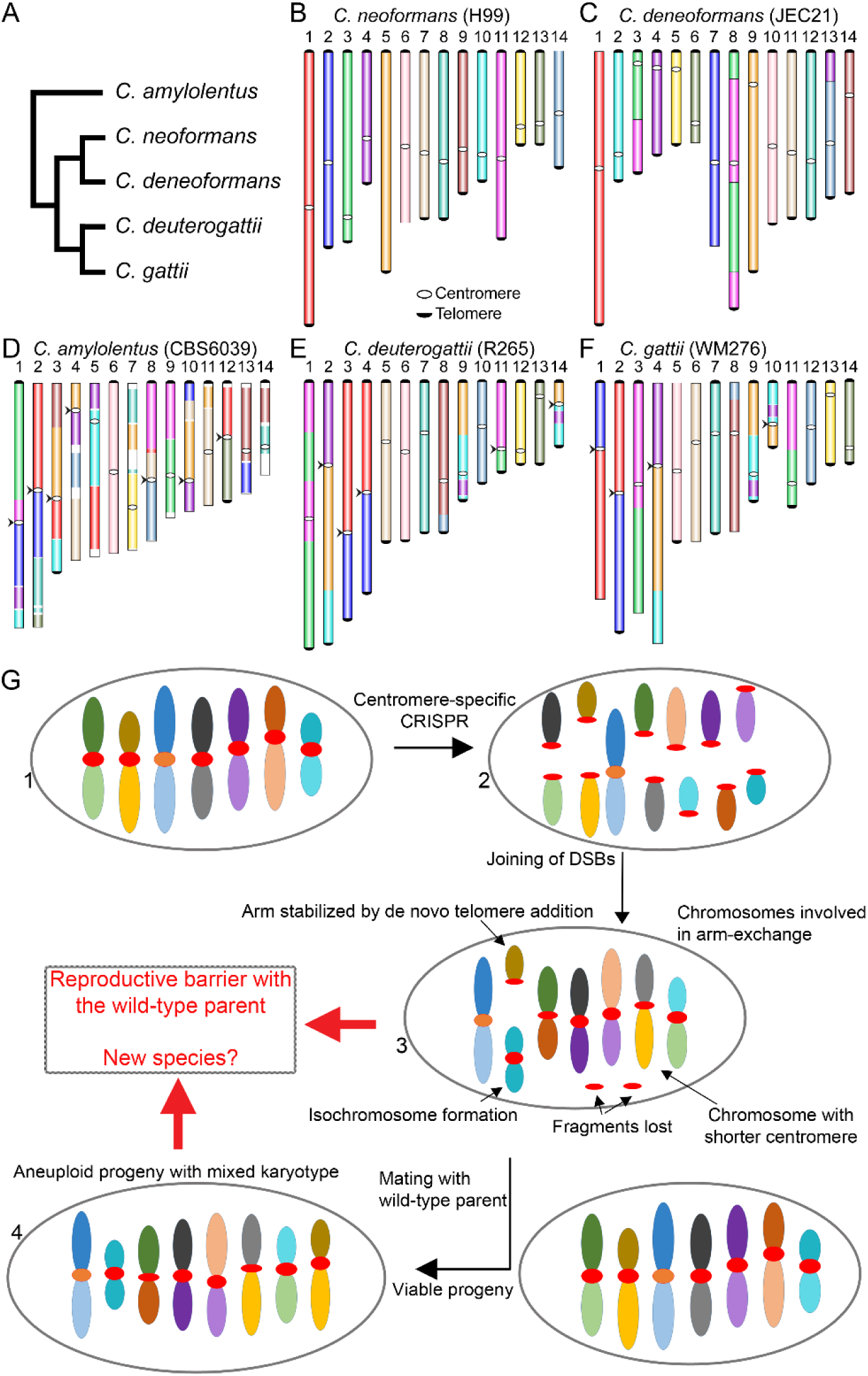
A model proposing the evolution of reproductive isolation induced by centromere breaks. **(A)** A schematic representing phylogeny showing the relationships of five *Cryptococcus* species, for which chromosome-level genome assemblies are currently available and published. **(B-F)** Chromosome maps for five species: *C. neoformans*, *C. deneoformans*, *C. amylolentus*, *C. deuterogattii*, and *C. gattii*. The synteny maps for all species were generated with *C. neoformans* as the reference and are colored accordingly. Chromosomal translocations involving centromeres are marked with arrowheads. **(G)** A model showing the events observed in the study. DSBs generated using CRISPR at centromeres (step 2) reshapes the karyotype following complex repair events. These complex events include the loss of centromere DNA, isochromosome formation, and *de novo* telomere formation (step 3), similar to what is observed during the process of chromothripsis. The new karyotype can generate a reproductive barrier with the parental isolate and lead to speciation. On the other hand, the strain with the rearranged karyotype could mate with wild-type isolate, albeit at low frequency, leading to aneuploid progeny (step 4), which can independently establish itself as a new species.

Recent studies have suggested that centromeres can undergo recombination, at least mitotic recombination, as compared to previous models in which centromeres were recalcitrant to recombination (22, 35–40). Our results further support that centromeres can undergo recombination when DSBs are generated in these regions. Each of the broken ends was processed and subjected to HR or NHEJ mediated repair and recombination. We also observed a case where a centromere, following the generation of DSBs, recombined, causing the loss of intermediate DNA sequences present between the two ends. In similar events, many fragments of centromere sequences were either lost or fused within other centromeres, altering the architecture of centromeres in this species (Fig. 7G). This result suggests that variation in centromere structure is not critical for centromere function, but might play some other role in genome organization. This conclusion is also supported by a previous study that showed a significant reduction in centromere length in *C. deuterogattii*, a closely related species of *C. neoformans* (22). This variation in centromere architecture also raises questions about the requirement of longer centromeres in most organisms. According to previously proposed models, longer centromeres with more centromere repeats might bind more kinetochore proteins (41–43). This would enable stronger kinetochore-microtubule interactions for longer centromeres and favor transmission of longer centromeres during meiosis. Our system can be harnessed to understand the dynamics of centromere transmission and test this hypothesis in future studies.

Interestingly, we also observed the addition of *de novo* telomere repeat sequences adjacent to the broken centromeres. While the mechanism underlying this process remains to be elucidated, the frequency of occurrence of *de novo* telomere addition seems to be high at ∼10% (3/32 ends). Additionally, two of the three telomere repeat additions involved invasion or fusion with other sequences before the addition of repeat sequences. *De novo* telomere formation on broken DNA ends has been observed in many species and is commonly known as chromosome healing (44–46). Chromosome healing occurs during mammalian development and has been observed in mouse embryonic stem cells as well as human germline cells (47–49). It is also proposed to occur in cancer cells to stabilize the ends of broken chromosome fragments arising as a result of chromothripsis or telomere crisis (50, 51). This process maintains the genome content while increasing the number of chromosomes in the case of DSBs that could not be repaired. *De novo* telomere addition in *S. cerevisiae* was shown to be influenced by the presence of telomere-like sequences at the broken DNA end. We found a similar scenario only in one of the three *de novo* telomeres formed, suggesting that the mechanism might differ in *C. neoformans*. In our study, we did not observe a significant impact of the loss of centromere sequences on the growth of the chromosomal shuffled strains as compared to the parental strain. This could suggest that shorter centromere sequences are sufficient to propagate the genome content. Furthermore, *de novo* telomere formation next to centromeres did not seem to have an effect on their functions suggesting that centromeres and telomeres do not influence each other’s function in *C. neoformans*, similar to telocentric/acrocentric chromosomes found in other organisms.

Repair of a DSB site is a complex process and involves multiple repair machineries including HR and NHEJ. HR mainly takes place during S-phase, whereas NHEJ occurs throughout the cell cycle (52, 53). HR mainly leads to gene conversion and can also drive recombination between repetitive sequences leading to translocation. NHEJ is more error-prone and can join any two sites resulting in translocations. Thus, both of these processes can lead to translocations but can result in very different types of junctions (8, 54). Our analysis of junctions exhibited evidence for both HR and NHEJ. Previous reports suggested that the NHEJ pathway is preferred over HR in the case of multiple DSBs (14, 15). Although the number is small, our results suggest that both of these pathways might take place at a similar rate in *C. neoformans*. However, the underlying regulatory mechanisms remain a subject for further investigation.

We also observed the inversion of sequences suggesting either strand invasion or simple fusion of a broken piece in reverse order by NHEJ. Our analysis also suggests the occurrence of multi-invasion repair (MIR), a subtype of HR, which can be favored for the repair of repeat sequences (6, 55). According to the MIR pathway, a single broken end might enter multiple target DNA molecules based on homology (56, 57). The DNA breaks in our experiments were made in the Tcn2 element, which is present in multiple copies across centromeres; thus, it is possible that some of the ends might have been repaired via MIR. Our results also suggest that breaks in repeat molecules are repaired either by NHEJ or HR, where other repeat sequences present in the genome may aid in the repair process. Similar observations have also been made previously in *S. cerevisiae*, where DSBs were generated with gamma radiation, HO-endonuclease, or CRISPR (58–60). In all of these studies, multiple copies of Ty elements were found to be involved in the generation of translocations. In the study that employed CRISPR, similar multiple translocation events were observed (59). However, Ty elements in *S. cerevisiae* are distributed across the genome, unlike *C. neoformans*, where all Tcn2 elements are clustered in centromeres.

Multiple concurrent breaks in the genome are rare and occur mainly in pathogenic conditions such as cancer. A commonly observed outcome is chromothripsis, where one or several chromosomes are broken into multiple smaller pieces and rejoined randomly to shuffle the targeted chromosomal region (11, 13). This phenomenon is associated with the generation of multiple mutations, as well as the activation of oncogenes. Chromothripsis is initiated by the generation of multiple breaks in a localized manner. The source of these breaks could be either internal factors such as replication associated breaks or external factors, including ionizing radiation or chemotherapy. These break sites are then repaired in a random manner. While this process is well-known, the mechanisms involved in this process are poorly understood (13, 61). In our experiments, we observed that similar events take place during DSBs repair (Step 3 in Fig. 7G). The connection between these processes needs to be further explored to establish whether our system can be extended to understand chromothripsis.

Our approach induced multiple simultaneous breaks in the genome, which were then repaired to generate chromosome shuffling. Centromeres are known to cluster in *C. neoformans* during mitosis (62, 63), and this may have promoted their interaction during the DSB repair events that generated these alterations. This centromere clustering in *C. neoformans* also mimics the clustering of DSB sites observed during the process of chromothripsis. We posit our approach could provide answers to critical questions regarding chromothripsis. The MIR pathway has been implicated as one pathway contributing to chromothripsis (6, 57). In our study, we also found evidence that the MIR pathway contributes to chromosome shuffling, further suggesting similarities between chromothripsis and the events we observed. Notably, chromothripsis is mainly observed in chromosome arms, whereas centromeres were targeted in our studies (13). However, some studies have indicated an association between chromothripsis break sites and the presence of transposon sequences (11, 64, 65). In our approach, the breaks were also located within transposon sequences, which are part of the centromeres. Thus, understanding the factors governing this process in *C. neoformans* could also shed light on facets of chromothripsis.

Chromosome rearrangements have been implicated in speciation, acting via reducing fertility or gene flow in the progeny that inherit the translocation (66–69). However, most models proposing speciation in this manner also consider geographical isolation as other major criteria (16). Our results, along with other studies, support this line of thought and suggest that chromosome rearrangements followed by geographical isolation can drive speciation (66, 70–72). Other models consider chromosome rearrangement as an effect of speciation, which happens as a result of simultaneous rearrangement after the speciation (67). A number of studies linking chromosome rearrangements and speciation have been conducted using *S. cerevisiae* and *Schizosaccharomyces pombe*. The *S. cerevisiae sensu stricto* group has been the subject of such studies over a long time period (73). A common approach has been to isolate viable hybrids between these species and then to study sporulation of these hybrids. These studies have suggested a role for three factors in the poor spore viability of hybrids: genetic incompatibilities, sequence divergence affecting mismatch repair system, and chromosome rearrangements (71, 74–76). However, more recent studies have suggested a greater role of chromosome translocations in this process than previously thought (77, 78). Studies involving a large collection of *S. pombe* isolates revealed karyotypic diversity among these isolates (79, 80). Some of these translocation events were found in geographically separated populations, suggesting a role in speciation. Another study found that *S. pombe* can mate successfully with a close relative, *Schizosaccharomyces kambucha* to form viable hybrids but these hybrids give rise to very few viable spores as compared to the individual parental species (81). The loss in viability was found to be correlated with two translocations present among these strains as well as three independent meiotic drive elements. Studies in marsupials suggested a role of chromosome rearrangements involving centromeres in the process of speciation in this species complex (82–86). These involved centromere-mediated translocations as well as differences in centromere lengths between two species. In our study, no genes were found to be affected or mutated because all rearrangements were generated via centromere recombination. Thus, a meiotic failure of rearrangement harboring strains suggests that chromosome translocations solely can drive reproductive isolation in a species. It is important to note that centromeres as such may not have any role in this phenomenon and rather centromere-localized breaks are just one of the ways to achieve chromosome rearrangements that in turn facilitate reproductive isolation. A recent study has provided evidence that even small repositioning of centromeres can generate reproductive barriers (87).

Here, we find that the presence of multiple centromere-mediated chromosomal rearrangements dramatically reduces the efficiency of meiosis and generates post-zygotic reproductive isolation. A complete failure to undergo meiosis with the existing population could lead to loss of new chromosome configurations, as they will not be passed to the new generations. We found that the presence of a mix of the original karyotype along with the new karyotype enhanced meiotic success. Interestingly, all three viable progeny exhibited similar karyotypes suggesting that this chromosome configuration is probably selected for cell survival while also keeping most of the genome as haploid. Thus, aneuploidy could also shield the newly established chromosome rearrangements and allow it to persist and spread in the population, eventually leading to the fixation of new changes. This hypothesis is supported by our results in which the aneuploid F1 progeny exhibited a much higher spore germination rate with either parent as compared to the crosses between the haploid parents. Additionally, the less aneuploid progeny exhibited less successful meiosis, further suggesting the role of aneuploidy in this process. Similar observations have also been made in *S. cerevisiae*, and its related species, where backcrosses and aneuploidy overcame reproductive isolation to increase spore germination (74, 75). Based on this, we propose a model in which the rearranged chromosomes can be present along with the parent chromosomes in an aneuploid intermediate (Fig. 7G). The aneuploid intermediate then allows the transmission of rearranged chromosomes to the next generation until a particular rearrangement becomes fixed in part of the population. Once fixed, the rearranged population will stably transmit itself and can give rise to a new subpopulation. The presence of geographical barriers, as well as possible advantageous selective roles for the new rearrangement, might further promote fixation of this newly acquired rearrangement.

## Acknowledgments

We thank Valerie Lapham at North Carolina State University electron microscopy facility for assistance with scanning electron microscopy. We thank Tom Petes and Sue Jinks-Robertson for critical reading of this manuscript. This work was supported by NIH/NIAID R37 MERIT award AI39115-21 to JH, R01 grant AI50113-15 to JH, and R01 grant AI112595-04 awarded to JH, David Tobin, and Paul Magwene. We also thank the CIFAR program, Fungal Kingdom: Threats & Opportunities for their support.

## Materials and Methods

### Strains and media

*C. neoformans* wild-type strains H99α and KN99**a** served as the wild-type isogenic parental lineages for the experiments. Strains were grown in YPD media for all experiments at 30°C unless stated otherwise, and G418 was added at a final concentration of 200 µg/ml for selection of transformants. MS media was used for all the mating assays, which were performed as described previously (26). Basidia specific spore dissections were performed after two weeks of mating, and the spore germination rate was scored after four days of dissection. All strains used in this study are listed in Table S3.

### Genome assembly and synteny comparison

The *C. neoformans* H99 genome was reassembled with Canu v1.7 based on previously published PacBio and Illumina data to obtain a better resolution of the centromeric regions (see Table S4 and S5 for details) (21, 22, 88). The resulting draft assembly was improved by correcting errors using five rounds of Pilon (v1.22) polishing (‘--fix all’ setting) and the Illumina reads mapped to the respective assemblies by the use of BWA-MEM (v0.7.17-r1188) (89, 90). Centromere locations were mapped based on BLAST analysis with centromere flanking genes, and coordinates for these new locations are provided in Table S1. Because some of the centromere lengths differed in the new assembly as compared to the previous one, we validated the new centromere lengths by mapping the Canu-corrected PacBio read using Minimap2, followed by visual inspection in IGV (91). Because our work involved analysis of centromere sequences, we utilized this new improved assembly as the reference for all of our analyses. Once the locations were mapped, we oriented all of the chromosomes such that the longer arm (q arm) begins the chromosome, and the smaller arm (p arm) is the distal part of the chromosome.

*De novo* genome assemblies for the VYD135 and VYD136 isolates, and progeny (P1, P2, and P3) were generated with Canu using Oxford Nanopore reads > 2 kb (see Table S4 and S5 for details) followed by Pilon polishing as described above. When necessary, and after validating by chromoblot analysis, broken contigs were joined artificially with a 50 bp sequence gap to generate a full-length chromosome. Once completed, the chromosomes were numbered based on their length with the longest chromosome as the first. For progeny P1, P2, and P3, extra chromosome configuration was inferred to obtain the final karyotype based on their Illumina and nanopore sequencing reads mapping. Sequence data generated in this study were submitted to NCBI with the BioProject accession number PRJNA577944.

Synteny comparisons between the genomes were performed with SyMAP v4.2 and default parameters (92) (http://www.agcol.arizona.edu/software/symap/). The comparison block maps were exported as .svg files and were then processed using Adobe Illustrator and Adobe Photoshop for representation purposes.

