## Supplementary material for "Centromere scission drives chromosome shuffling and reproductive isolation": Suplementary Information

This PDF file includes:

SI Materials and Methods

Figures S1 to S8

Tables S1 to S5

SI References

### SI Materials and Methods

#### *Phenotyping and growth competition assays*

For cell growth and phenotyping assays, cultures were grown overnight, serially diluted (10-fold dilutions), and 3 µl of cell suspension was spotted on YPD or defined stress solid medium followed by incubation at 30°C and 37°C for two days. To make drug-containing media, fluconazole and phleomycin were added into YPD media at a concentration of 24 µg/ml and 1 µg/ml, respectively, and plates were made. The plates were imaged after two days of growth and are presented.

To test the competitive fitness, H99, VYD135, and VYD136 strains were separately mixed with CNVY156 (1) at an equal ratio based on OD measurements. CNVY156 strain expresses NAT resistant marker and was used as the counter-selective strain to score competitive growth in co-culture experiment. After the strains were mixed, a part of the mix (1 ml) was immediately spread on YPD plates (0h sample), and cells were allowed to grow for 48 h at 30°C. The remaining mix (5 ml) was grown as co-culture at 30°C for 24 h, and then a portion of cells from the mix was again spread on YPD plates (24h sample) and incubated for 48 h. From both initial and final samples, 100-150 randomly selected single colonies were replica patched on YPD and YPD+NAT media plates. After 24 h of growth, number of growing colonies from both YPD and YPD+NAT plates was counted. The growth fraction for H99, VYD135 and VYD136 strains was calculated by subtracting number of colonies on YPD+NAT plate from the total number of colonies on YPD plate for each co-culture. The percent growth was calculated and plotted using GraphPad with the total number of colonies on YPD representing 100%, the number of NAT resistant colonies representing $NAT^{+}$ population while remaining colonies were considered to be either H99, VYD135, and VYD136.

#### *CRISPR transformations*

CRISPR transformation experiments were performed as described previously, with minor modifications (2) (Fig. S1). Briefly, *C. neoformans* cells were grown overnight in YPD media and then inoculated into 100 ml of fresh YPD media and grown until $OD_{600}$ reached 0.8 to 1. The cells were then pelleted, washed, and resuspended in EB buffer and incubated for one hour. Next, the cells were pelleted and resuspended again in 50 µl of EB buffer. The cells were mixed with three DNA fragments expressing guide RNA (350 ng), Cas9 (500 ng), and selection marker (2 µg). The cell-DNA mix was subjected to electroporation using the Eppendorf multiporator, with the bacterial mode operating at V=2 kV with τ optimized at 5 ms. Fresh YPD media was immediately added into the transformed cells, and the cells were allowed to recover for two hours before spreading onto selection media (YPD+200 µg/ml G418). The transformants were recovered after 3 to 5 days after transformation. The transformants were further colony purified on YPD+G418 medium to obtain single colony stable transformants. The guide RNA coding sequences used for Tcn2 and safe-haven locus targeting are TAAGTACTTCTGGGATGGTA and AGTGCTGTGGTGAAAGAGAT, respectively.

#### *PFGE and chromoblot analysis*

The PFGE plugs were prepared as described previously with minor modifications (3). The PFGE was performed with 1% agarose gel at 3.6 V/cm and a switching frequency of 120 to 360 sec for 120 h at 14°C in 0.5X TBE. *S. cerevisiae* CHEF DNA marker (Bio-rad, Cat #1703605) served as markers for estimating the chromosome lengths in all PFGE experiments. To separate shorter chromosomes, the switching frequency was changed to 116 to 276 sec, while all other conditions were kept the same. Following electrophoresis, gels were stained with ethidium bromide (EtBr), and bands were observed by UV transillumination and photographed. Chromoblot analysis was performed as described previously (4). Briefly, the DNA was hybridized to membranes and probed with chromosome arm regions from targeted chromosomes. The probed membranes were washed, and hybridization signals were observed with a phosphorimager.

***Genomic DNA isolation for nanopore sequencing***

The strains with chromosome alterations, as well as progeny from the VYD136α x KN99**a** cross, were subjected to nanopore and Illumina sequencing. For nanopore sequencing, large molecular weight genomic DNA was prepared using the CTAB based lysis method. For this purpose, 50 ml of an overnight culture was pelleted, frozen at -80°C, and subjected to lyophilization. The lyophilized cell pellet was broken into a fine powder by vortexing with beads for 3 to 5 min at room temperature. 20 ml of CTAB extraction buffer (100 mM Tris-Cl, pH=7.5; 0.7 M NaCl; 10 mM EDTA, pH=8.0; 1% CTAB; 1% β-mercaptoethanol) was added, mixed, and incubated at 65°C for an hour with intermittent shaking after every 20 min. The mix was cooled on ice for 10 min, and the supernatant solution was decanted into a fresh tube. An equal volume of chloroform (~15 ml) was added to the tube and mixed gently for 5 to 10 min.

The mix was centrifuged at 3200 rpm for 10 min, and the supernatant was transferred to a fresh tube. An equal volume of isopropanol (~18 to 20 ml) was added into the supernatant and mixed gently until thread-like structures were visible and formed a clump. The mix was incubated at -20°C for an hour, and centrifuged at 3200 rpm for 10 min to pellet the DNA. The supernatant was discarded, and the pellet was washed with 70% ethanol. The pellet was air-dried and dissolved in 1ml of 1X TE buffer. RNase A was added into the resuspended DNA to a final concentration of 100 µg/ml and incubated at 37°C for 30 to 45 min. Sodium acetate solution was added into the mix to a final concentration of 0.5 M, and the solution was transferred to a 1.5 ml Eppendorf tube/s in the aliquots of 0.5-0.6 ml each. An equal volume of chloroform was added, mixed gently, and centrifuged at 13,000 rpm for 15 min. The supernatant was transferred to a fresh tube, followed by isopropanol precipitation. The DNA threads were removed with a pipette tip and transferred to a fresh tube followed by ethanol washing, air drying, and finally dissolved in 200 µl 1X TE buffer. The DNA quality was estimated with NanoDrop whereas DNA quantity was estimated with Qubit. The size estimation of DNA was carried by electrophoresis of DNA samples on PFGE. For this purpose, the PFGE was carried out at 6V/cm at a switching frequency of 1 to 6 sec for 16 h at 14°C. Samples with most of the DNA ≥100 kb or larger were selected for nanopore sequencing.

***Nanopore and Illumina sequencing***

The DNA samples isolated as above were subjected to library preparation and sequencing, as recommended by the manufacturer's instructions. For nanopore sequencing, three or four different DNA samples were barcoded as per the manufacturer's instructions using SQK-LSK109 and EXP-NBD103/EXP-NBD104 kits. The DNA samples were then pooled together on a single R9 flow-cell (FLO-MN106), and sequenced by the MinION system. MinION sequencing and live-base calling were controlled using MinKNOW. DNA sequencing was performed at the default voltage for 48 hours. After sequencing, reads were de-multiplexed with qcat (https://github.com/nanoporetech/qcat) (parameters: --trim -k NBD103/NBD104 --epi2me). Each set of reads was then assembled to obtain genome assembly using Canu as described previously.

Illumina sequencing of the strains was performed at the Duke sequencing facility core (https://genome.duke.edu/), and the data was employed to error correct the genome assemblies for VYD135 and VYD136. The Illumina sequencing data were also mapped to the wild-type H99 genome assembly to estimate the ploidy of strains. Specifically, the Illumina reads were mapped to the H99 genome using Geneious default mapper or bow-tie2 mapper. The resulting BAM file was converted to a .tdf file, which was then visualized through IGV to estimate the ploidy based on read coverage for each chromosome.

To map the breakpoints, nanopore reads from both VYD135 and VYD136 were mapped to the wild-type genome assembly using minimap2. The mapped file (.bam) file was visualized in IGV, and centromere specific snapshots were exported for representation. Comparative analysis of the new centromere sequences with parental sequences was performed by both BLASTn analysis and pairwise sequence alignment. The *CAS9* and *NEO* gene sequences were identified in the new centromeres based on BLAST analysis with sequences of original plasmids.

***Galleria mellonella killing assay***

*G. mellonella* infection experiments were performed as described previously with some modifications (5). *G. mellonella* caterpillars in the final instar larval stage were used to test the pathogenicity of *C. neoformans* strains, and healthy caterpillars were employed in all assays. 20 to 22 chosen caterpillars were infected with each strain. Four µl cell suspension ($10^6$ cells/ml) of a strain was injected into each caterpillar via the last left proleg. After injection, caterpillars were incubated in plastic containers at 37°C, and the number of dead caterpillars was scored daily. Caterpillars were considered dead when they exhibited black coloration of the body and displayed no movement in response to touch. The few caterpillars that became pupae during the experimental duration were removed from the analysis.

### Supplementary Figure and Figure Legends

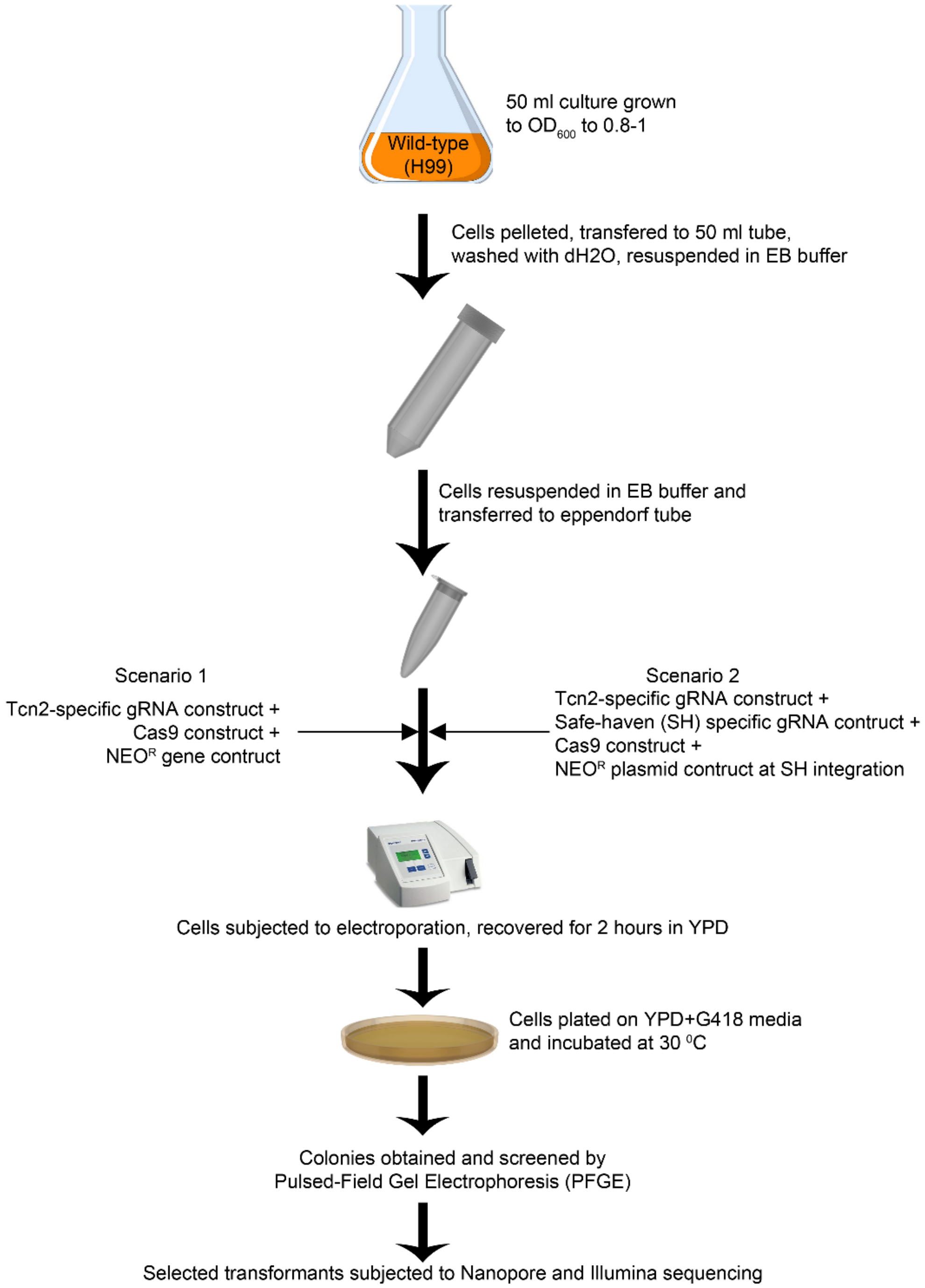


**Fig. S1. A schematic diagram illustrating the experimental procedure.** An outline detailing the protocol for CRISPR transformation is shown here. The cells were transformed with either one guide RNA, where the selection marker was targeted to the centromere, or two guide RNAs, where selection marker was targeted for integration at the safe haven locus instead of the centromere.

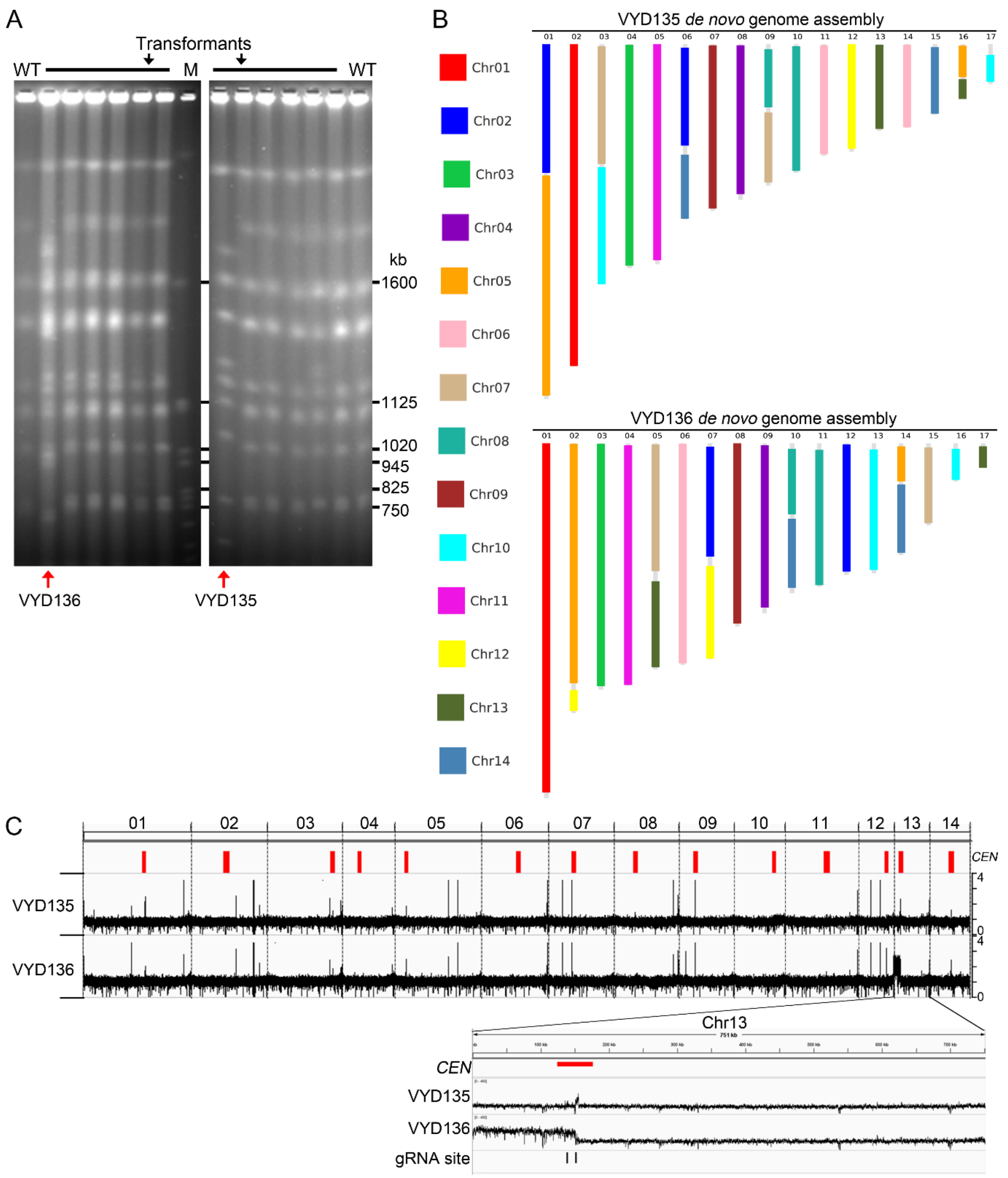


**Fig. S2. Nanopore and Illumina sequencing for chromosome shuffled strains, VYD135, and VYD136. (A)** EtBr stained PFGE showing chromosome band pattern for transformants. WT, wild-type H99; M, *S. cerevisiae*. **(B)** *De novo* genome assembly showing contigs for the VYD135 and VYD136 strains. The contigs are colored based on their synteny with wild-type chromosomes. While both the genome assemblies were free of any errors or sequence gaps, a region of chromosome 2 (next to the centromere), was found to be missing in VYD13. Illumina reads showed that this region is not deleted in VYD136 but is absent in the assembly due to an assembly error. **(C)** Illumina sequencing confirmed the haploid nature of VYD135 and VYD136, with the only exception being the smaller arm of chromosome 13 in VYD135, which is present in two copies.

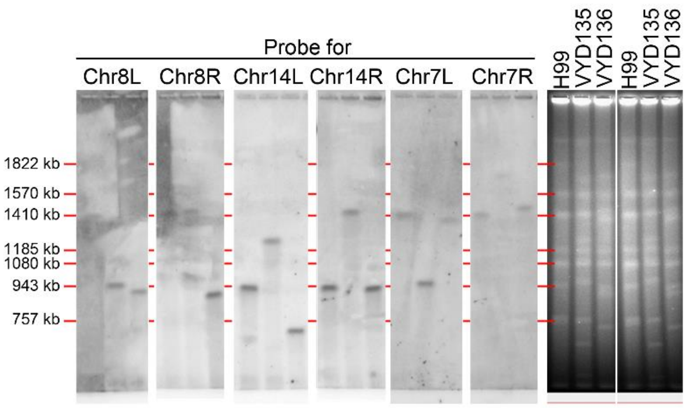


| | 8L | 8R | 14L | 14R | 7L | 7R |
|---|---|---|---|---|---|---|
| H99 | 1406 | 1406 | 943 | 943 | 1413 | 1413 |
| VYD135 | 1004 | 1412 | 1245 | 1412 | 1004 | 1704 |
| VYD136 | 967 | 926 | 724 | 967 | 1366 | 1457 |

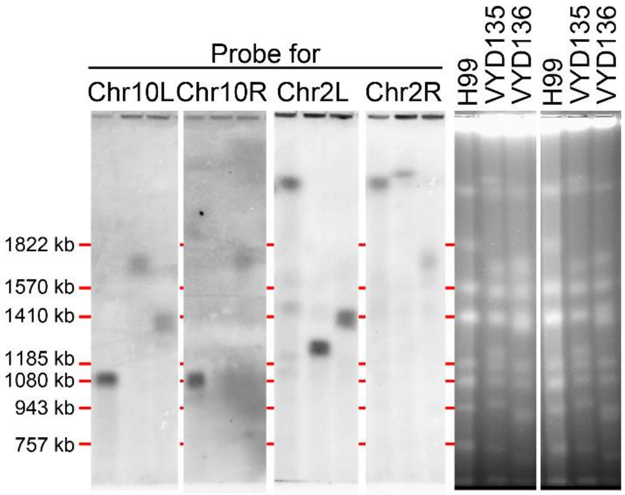


| | 10L | 10R | 2L | 2R |
|---|---|---|---|---|
| H99 | 1073 | 1073 | >2000 | >2000 |
| VYD135 | 1704 | 284 | 1245 | 2513 |
| VYD136 | 1366 | 1093 with rDNA | 1391 | 1093 with rDNA |

**Fig. S3. Chromoblot analysis confirms chromosomal translocations in strains, VYD135, and VYD136.** L and R represent the left and right arms of the chromosome for which specific probes were hybridized. Green colored numbers in the table represent the length (in kb) of confirmed bands, whereas the black number represents the size of the unconfirmed band that could not be detected due to its small size.

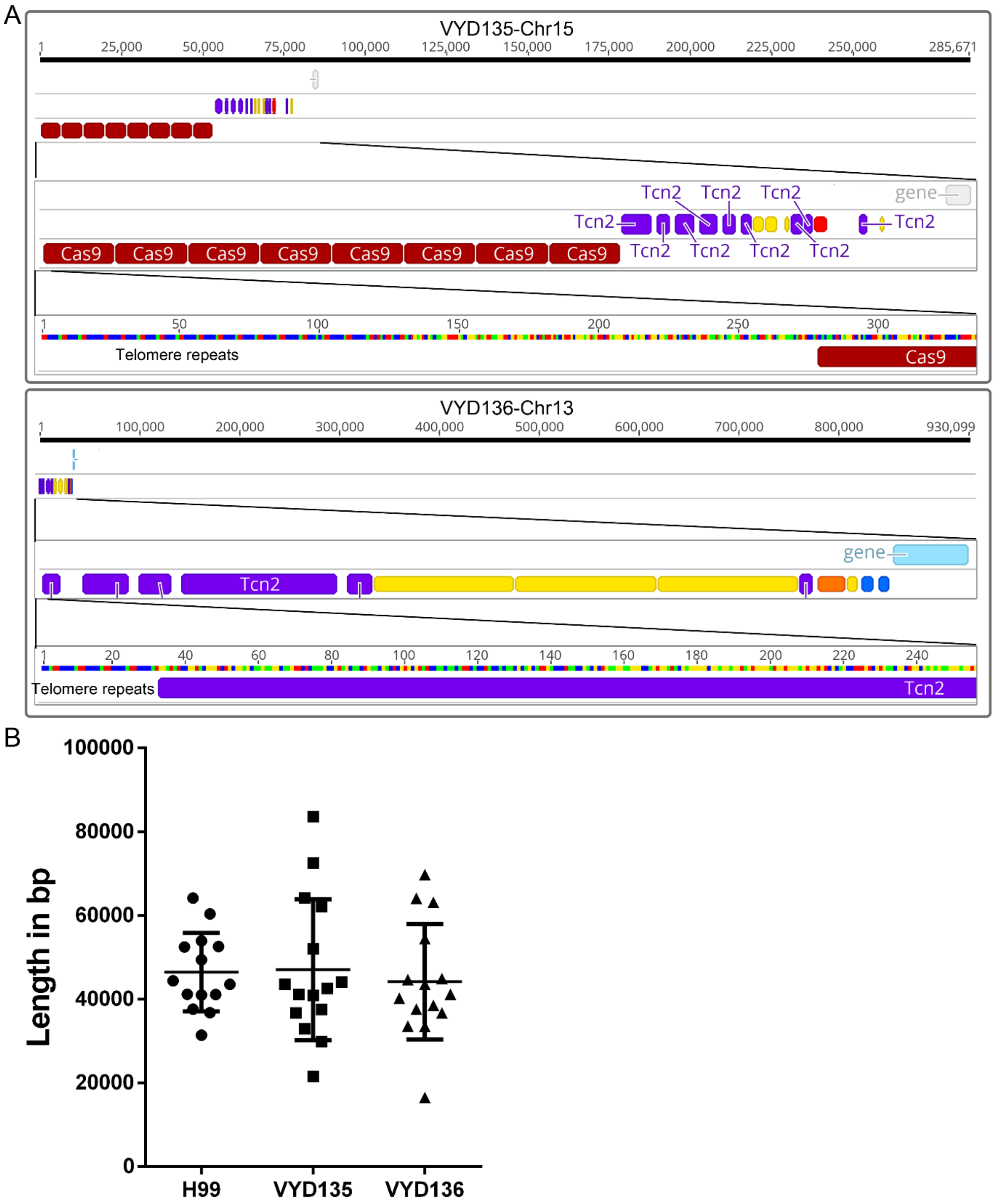


**Fig. S4. Chromosome rearrangements generated new telomeres and shortened centromeres. (A)** Two new telomeres that formed on chromosome 15 of VYD135 and chromosome 13 of VYD136 are shown here. In chromosome 15 of VYD135, the centromeric sequence has been fused with the Cas9 sequence, and this was then followed by the addition of telomere repeat sequences. **(B)** A graph showing the lengths of centromeres in H99, VYD135, and VYD136.

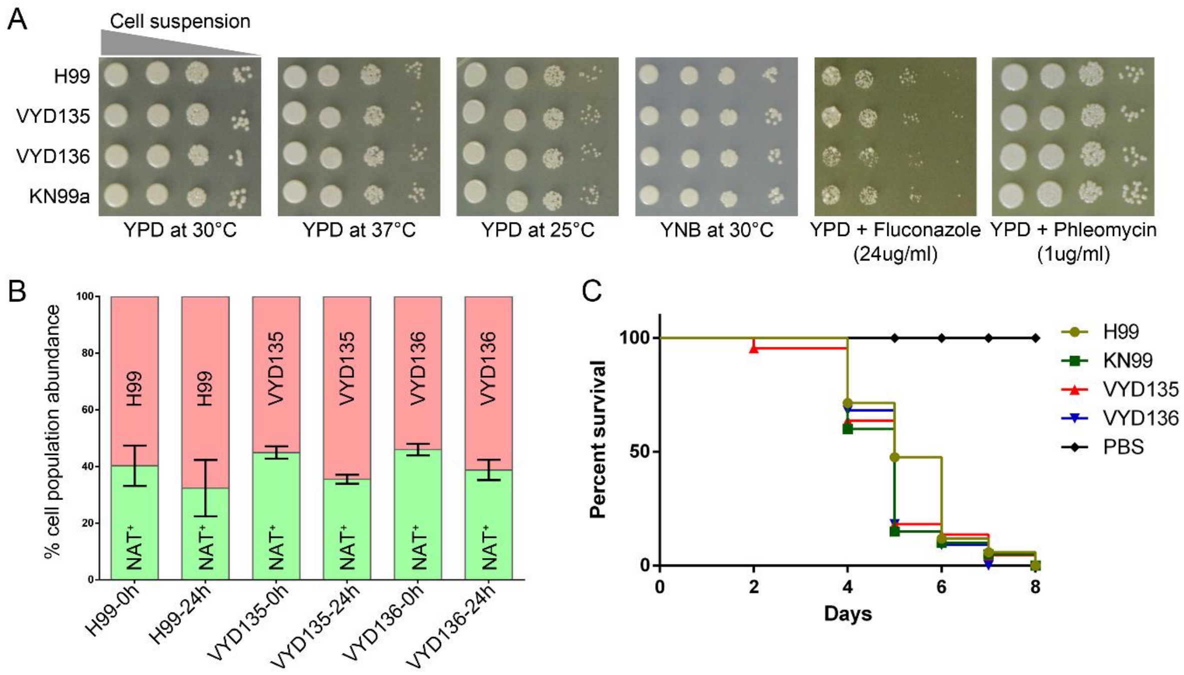


**Fig. S5. Growth and pathogenicity of VYD135 and VYD136 strains. (A)** A plate spotting assay showing the growth of wild-type (H99 and KN99**a**) and rearranged strains (VYD135 and VYD136) at different temperatures, nutrient limiting (YNB) and drug-containing (fluconazole and phleomycin) media conditions. **(B)** A graph showing the competitive growth fitness of H99, VYD135, and VYD136 strains in a co-culture experiment after 24 hours. **(C)** A survival plot showing the virulence of *C. neoformans* strains as compared to the PBS control. No significant differences were observed in the virulent phenotypes of the wild-type H99 or KN99**a** strains as compared to either VYD135 or VYD136 strains.

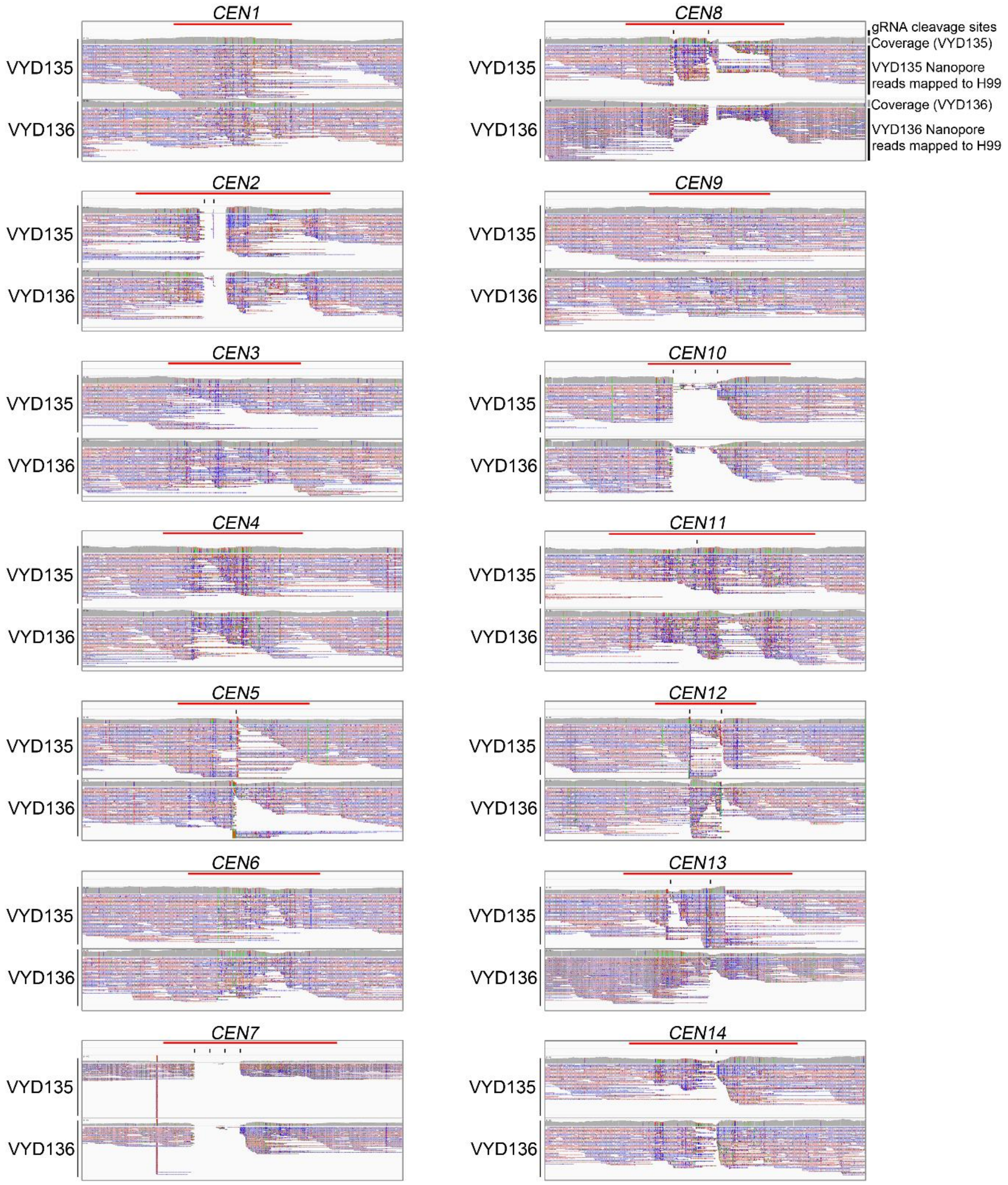


**Fig. S6. Analysis of nanopore reads mapped DSBs to gRNA cleavage sites.** A 100 kb region encompassing the centromeres is shown for each of the 14 centromeres in the H99 genome. Mapping of the VYD135 and VYD136 nanopore reads to these regions revealed that the DSB sites coincide with the gRNA target sites. The red bars represent the position of the centromeres, and black lines denote the sites of gRNA cleavage.

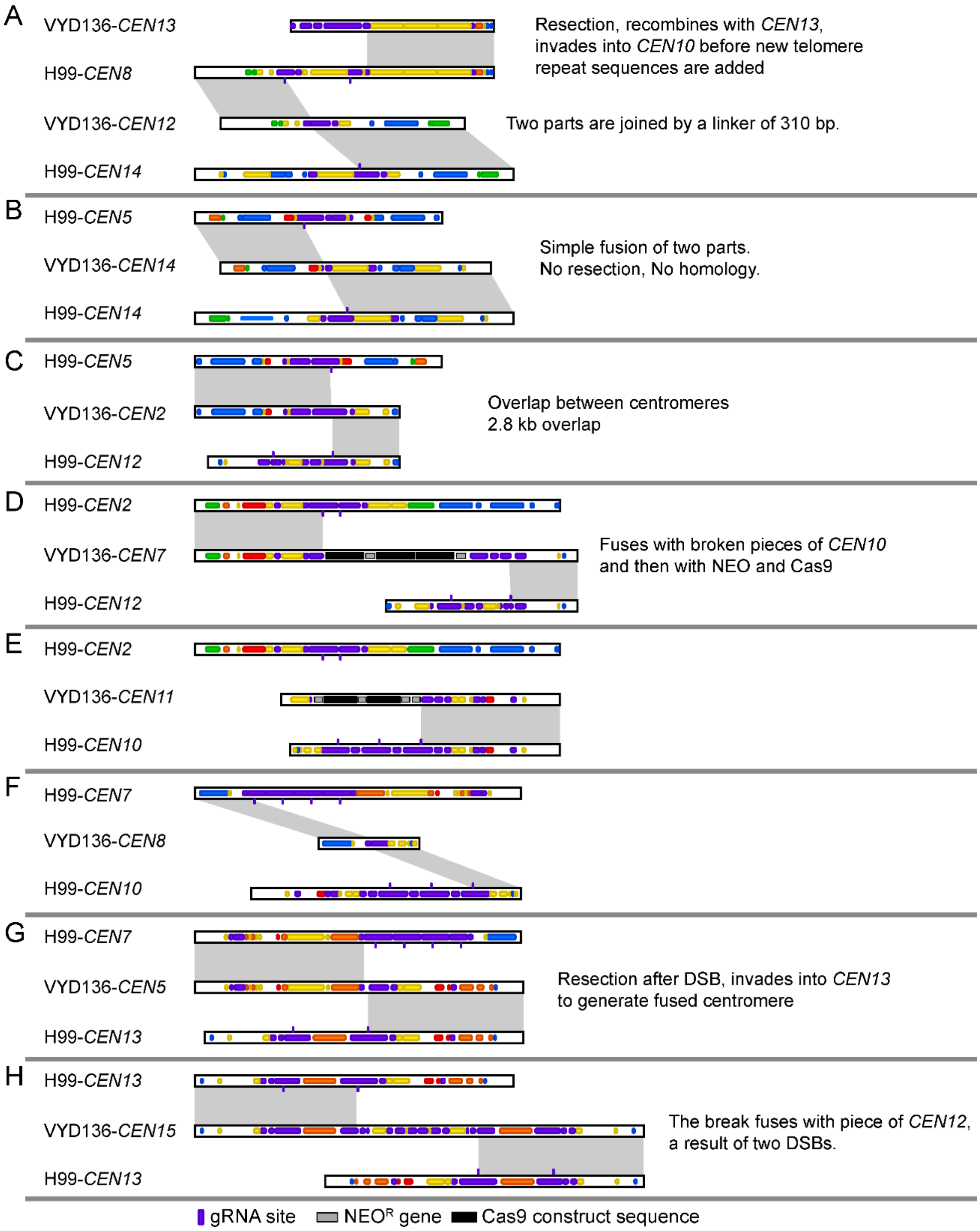


**Fig. S7. Synteny analysis of rearranged centromeres compared to native centromeres. (A-H)** Synteny comparison of VYD136 centromeres with wild-type H99 centromeres revealed multiple complex rearrangement events. The synteny between H99-*CEN2* and VYD136-*CEN11* (E) could not be mapped due to a break in the VYD136 genome assembly. Grey shades mark direct synteny between the two sequences.

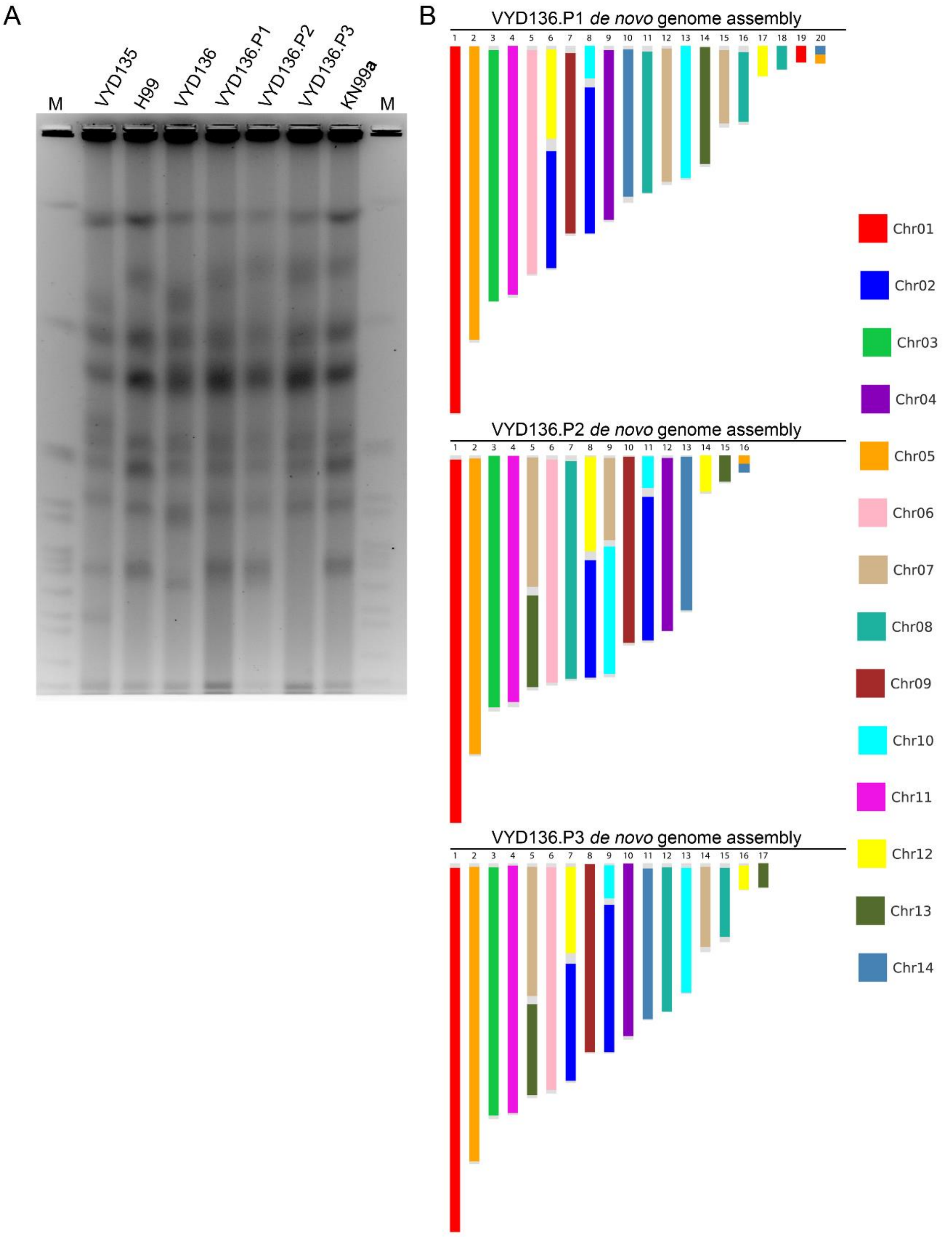


**Fig. S8. *De novo* genome assembly for VYD136 progeny following nanopore sequencing.** **(A)** An EtBr stained PFGE gel image showing the chromosome bands for VYD136 x KN99**a** cross progeny, P1, P2, and P3. **(B)** *De novo* genome assembly showing contigs for the progeny obtained from the VYD136 X KN99**a** cross. The contigs are colored based on their synteny with wild-type chromosomes.

### Supplementary Tables

**Table S1.** ***De novo*** **telomere repeat sequences identified in VYD135 and VYD136.**

| **Chromosome number** | **Sequence** |
|---|---|
| VYD135-Chr13 | ACCAAGTGCCCCACTCAGACGCGCACTTCGACACCCAAAGTCCATCATT<br>TGGAGACCGAACTGACGGCGGGTGACGCTACCTCGGGGTATCTCACGTT<br>AGCCCAGACCCTTCCTCCAGCCGATGATGGCAACGGTGATGCTACGTAC<br>GGG<u>TTAGGGGTTAGGGGTTAGGGGTTAGGGGTTAGGGGTTAGGGGTTA</u><br><u>GGGGTTAGGGGTTAGGGGTTAGGGGTTAGGGGTTAGGGGTTAGGGGTT</u><br><u>AGGGGTTAG</u> |
| VYD135-Chr15 | AAACCAGTCATTGAACGGTAACAAGCATAACTTCCGATGTAGCTTACCG<br>CCTGCCACTCGTCCCACTGTTCATCACCCGCTTCGATCTTATGCTCAACC<br>TCGTTCCAATTCATACCCGACTTTTCCAAGACGTCACTGATTGGATTGAT<br>G<u>TTAGGGGGTTAGGGGGTTAGAGGGTTAGGGGGTTAGGGGGTTAGGGG</u><br><u>TTAGGGGTTAGGGGTTAGGGGTTAGGGGTTAGGGGTTAGGGGTTGGGG</u><br><u>TTGGGGGGTTAGGGGTTAGGGGTT</u> |
| VYD136-Chr13 | CCCCCACGATACCTCAGGCCCAACCAACTGCGACGGAATGGGACTTAG<br>CAGCGAAGAGGATGGAGTTAGAAGAAGGTATCGCTCGTGACGAACTGA<br>AGCATCGGCAAAGTGTTCAAGCCAACAAGCACCGCTGTCCCGATCCGG<br>TATACC<u>TAGGGGTTAGGGGTTAGGGGTTAGGGGTTAGG</u> |

Sequences for de novo telomeres along with 150-bp of preceding sequences are shown for the three newly formed telomeres. The telomere repeat ($TTAG_{4-5}$) sequences are marked as underlined sequence.

**Table S2. Centromere locations in strains H99, VYD135, and VYD136.**

| *CEN* (Chr) # | H99 | VYD135 | VYD136 |
|---|---|---|---|
| *CEN1* | 1288448-1325216 (36769) | 1572354-1634513 (62160) | 1288245-1324985 (36741) |
| *CEN2* | 847968-908364 (60397) | 1287601-1324342 (36742) | 1572350-1605934 (33585) |
| *CEN3* | 1373375-1417998 (44624) | 828082-880128 (52047) | 1372843-1417426 (44584) |
| *CEN4* | 729460-773021 (43562) | 1372860-1417465 (44606) | 874657-938791 (64135) |
| *CEN5* | 1575333-1616363 (41031) | 874656-938807 (64152) | 828964-883407 (54444) |
| *CEN6* | 782846-823959 (41114) | 779176-820076 (40901) | 782697-821224 (38528) |
| *CEN7* | 829116-883060 (53945) | 896582-940727 (44146) | 730008-793154 (63147) |
| *CEN8* | 896639-946048 (49410) | 730015-802561 (72547) | 831898-848437 (16540) |
| *CEN9* | 806454-844077 (37624) | 806365-843963 (37599) | 806317-843923 (37607) |
| *CEN10* | 832012-876442 (44431) | 714887-758436 (43550) | 714910-758463 (43554) |
| *CEN11* | 874913-939101 (64189) | 525138-555048 (29911) | 851610-898623 (47014) |
| *CEN12* | 604242-635674 (31433) | 603993-625596 (21604) | 453574-493788 (40215) |
| *CEN13* | 579568-632178 (52611) | 577624-620105 (42482) | 896597-930068 (33472) |
| *CEN14* | 490223-542717 (52495) | 235247-268182 (32936) | 447549-492438 (44890) |
| *CEN15* | - | 201940-285540 (83601) | 124270-198037 (73768) |

**Table S3. Strains used in this study.**

| Strain name | Description | Reference |
|---|---|---|
| H99 | Wild-type α | (6) |
| KN99**a** | Wild-type **a** | (7) |
| VYD135 | H99 derivative with shuffled chromosomes | This study |
| VYD136 | H99 derivative with shuffled chromosomes | This study |
| VYD146 | Progeny from the cross between VYD136 and KN99**a** (VYD136.P1) | This study |
| VYD147 | Progeny from the cross between VYD136 and KN99**a** (VYD136.P2) | This study |
| VYD148 | Progeny from the cross between VYD136 and KN99**a** (VYD136.P3) | This study |
| CNVY156 | GFP-CENP-A-NAT$^{R}$ expressing strain | (1) |

### Table S4. PacBio, Nanopore, and Illumina read data used in this study.

| Raw data files | Strains | Sequencing Technology | Number of reads | Total bases (bp) | Read Length (bp) | Mean read length (bp) | Read length N50 (bp) | Max. read length (bp) | GenBank Acc. No. | Purpose |
|---|---|---|---|---|---|---|---|---|---|---|
| **PacBio reads** | | | | | | | | | | |
| **H99_0doubling_subreads.10k.fastq** | H99 | PacBio, Sequel | 46,202 | 675,769,783 | | 14,626 | 14,428 | 56,652 | SRR6435868 | *de novo* assembly |
| **H99_1000doubling_subreads.fastq** | | PacBio, Sequel | 834,406 | 8,346,702,363 | | 10,003 | 15,768 | 100,682 | SRR6435871 | |
| **Nanopore reads** | | | | | | | | | | |
| **VYD135.barcode02.run1.fastq** | VYD135 | ONT, MinION | 288,634 | 1,296,872,891 | | 4,493 | 16,191 | 156,671 | SRR10317001 | *de novo* assembly |
| **VYD135.barcode02.run3.fastq** | VYD135 | ONT, MinION | 112,421 | 522,271,300 | | 4,645.67 | 12,726 | 130,456 | | |
| **VYD136.barcode03.run1.fastq** | VYD136 | ONT, MinION | 314,779 | 1,178,878,974 | | 3,745 | 12,187 | 179,586 | SRR10317000 | *de novo* assembly |
| **VYD136.barcode03.run3.fastq** | VYD136 | ONT, MinION | 69,796 | 400,748,478 | | 5,741 | 15,376 | 224,370 | | |
| **VYD136.P1.barcode09.fastq** | VYD136.P1 | ONT, MinION | 270,168 | 1,614,653,148 | | 5976.48 | 17,096 | 175,846 | SRR10316999 | *de novo* assembly |
| **VYD136.P2.barcode10.fastq** | VYD136.P2 | ONT, MinION | 1,254,515 | 4,614,488,350 | | 3678.3 | 10,206 | 218,170 | SRR10316998 | *de novo* assembly |
| **VYD136.P3.barcode11.fastq** | VYD136.P3 | ONT, MinION | 649,725 | 2,176,664,992 | | 3350.13 | 10,301 | 214,885 | SRR10316997 | *de novo* assembly |
| **Illumina reads** | | | | | | | | | | |
| **SRR642222_{1,2}c.fastq** | H99 | Hiseq 2000 | 7,757,786 | 783,536,386 | 101, PE | - | - | - | SRR642222 | Pilon correction of H99 assembly |
| **SRR647805_{1,2}.fastq** | H99 | Hiseq 2000 | 8,041,332 | 812,174,532 | 101, PE | - | - | - | SRR647805 | |
| **VYD135.fastq** | VYD135 | Hiseq 4000 | 11,187,790 | 3,378,712,580 | 151, PE | - | - | - | SRR10317030 | Pilon correction of VYD135 assembly |
| **VYD136.fastq** | VYD136 | Hiseq 4000 | 13,428,856 | 4,055,514,512 | 151, PE | - | - | - | SRR10317029 | Pilon correction of VYD136 assembly |
| **VYD136-P1.fastq** | VYD136.P1 | Hiseq 4000 | 7,424,959 | 2,242,337,618 | 151, PE | - | - | - | SRR10911080 | Pilon correction of VYD136.P1 assembly |
| **VYD136-P2.fastq** | VYD136.P2 | Hiseq 4000 | 6,522,372 | 1,969,756,344 | 151, PE | - | - | - | SRR10911079 | Pilon correction of VYD136.P2 assembly |
| **VYD136-P3.fastq** | VYD136.P3 | Hiseq 4000 | 6,532,368 | 1,972,775,136 | 151, PE | - | - | - | SRR10911078 | Pilon correction of VYD136.P3 assembly |

**Table S5. Statistics for genome assemblies generated in this study.**

| | H99 | VYD135 | VYD136 | VYD136.P1 | VYD136.P2 | VYD136.P3 |
|---|---|---|---|---|---|---|
| *De novo* Contigs | 14 | 17 | 17 | 20 | 16 | 17 |
| Final chromosomes | 14 | 15* | 15* | 16* | 17* | 15* |
| Total length | 19,089,897 | 19,125,078 | 19,045,361 | 19,454,758 | 19,299,987 | 19,088,103 |
| Telomeres | 26 | 30 | 30 | 33 | 28 | 27 |
| Accession number | PRJNA577944 (CP047902-CP047915) | PRJNA577944 (JAACNN000000000) | PRJNA577944 (JAACNM000000000) | PRJNA577944 (JAACNL000000000) | PRJNA577944 (JAACNK000000000) | PRJNA577944 (JAACNJ000000000) |

Accession number provided are for the *de novo* assemblies.

*The final genome assemblies included some chromosome configurations that were predicted based on PFGE and ploidy data obtained from nanopore and Illumina sequencing reads.
